# Defective lysosomal acidification contributes to TNFR1 mediated neuronal necroptosis in Alzheimer’s disease

**DOI:** 10.1101/2023.10.12.562041

**Authors:** Jialiu Zeng, Kholoud Abd-Elraouf, Gavin Wen Zhao Loi, Eka Norfaishanty Saipuljumri, Lance M. O’Connor, Jonathan Indajang, Richard Reynolds, Anna M. Barron, Chih Hung Lo

## Abstract

**Background:** Tumor necrosis factor (TNF) receptor 1 (TNFR1) signaling mediates neuronal necroptosis in Alzheimer’s disease (AD). Interaction of TNFR1 signaling axis with autolysosomal pathway and the accumulation of necrosome molecules in impaired lysosomes have been shown to lead to necroptotic neuronal death. This has been attributed to the terminal failure of the autophagic process, primarily due to lysosomal degradation dysfunction. Being the final and determining step of the autolysosomal pathway, lysosomes with sufficient acidification as maintained by functional vacuolar (H+)-ATPase (V-ATPase) are required to achieve complete autophagic degradation of toxic cellular components. Here, we aim to investigate the role of defective lysosomal acidification in mediating TNFR1 induced neuronal necroptosis in AD.

**Methods:** Neuropathological analysis of human post-mortem AD brains was performed to examine the correlation between TNFR1 induced neuronal necroptosis and autolysosomal dysfunction. Specifically, we probed for the level of V-ATPase subunits in AD brains to determine the extent of lysosomal acidification and function. Cell-based assays were conducted to understand the effect of TNFR1 activation in driving lysosomal acidification defect, proteolytic function, membrane integrity, autophagic impairment, mitochondrial dysfunction, and neuronal death in SH-SY5Y neuroblastoma cells. Furthermore, we applied lysosome-acidifying nanoparticles (AcNPs) to determine whether restoration of lysosomal acidification can rescue neuronal necroptosis in both TNF-treated SH-SY5Y cells and APP^NL-G-F^ knock-in mouse model of AD.

**Results:** We found that TNFR1 activated neuronal necroptosis correlated with autolysosomal dysfunction as characterized by downregulation of V-ATPase subunits and accumulation of autophagy receptor p62 in human AD brains. In cell culture, we showed for the first time that lysosomal acidification is only impaired in cells treated with TNF and not with other cytokines, contributing to inhibition of autophagic degradation in SH-SY5Y cells. TNF also disrupted lysosomal trafficking and membrane dynamics and induced lysosomal membrane permeabilization, followed by impaired autophagic clearance, defective mitochondrial turnover, reduced mitochondrial function, and neuronal death. Importantly, we demonstrated that AcNPs restored lysosomal, autophagic, and mitochondrial function, improved lysosomal membrane homeostasis, and rescued neuronal necroptosis in both TNF-treated SH-SY5Y cells and APP^NL-G-F^ mice.

**Conclusions:** Defective lysosomal acidification plays a key role in TNFR1 mediated neuronal necroptosis. This opens avenues for new therapeutic strategies to target lysosomal acidification dysfunction in AD.

## Background

Alzheimer’s disease (AD) is neurological disorder characterized by sustained neuroinflammation leading to progressive neurodegeneration with clinical symptoms of cognitive decline and memory loss [1]. Alongside the classical pathological hallmarks of intracellular tau neurofibrillary tangles (NFTs) and extracellular β-amyloid (Aβ) plaques, increasing evidence suggests that neuroinflammation plays a prominent role in AD pathogenesis [2,3]. This is supported by AD risk genes that are associated with immune response [4,5] and the elevated levels of inflammatory cytokines in AD patients [6]. It has also been shown that neuroinflammation actively drives AD pathogenesis, rather than a mere bystander or consequence of insults [2,3]. One of the key inflammatory mediators under neurodegenerative condition is the soluble tumor necrosis factor (TNF) that primarily acts through binding TNF receptor 1 (TNFR1) and activating downstream signaling, including cell death pathways [7]. There is also growing evidence that TNFR1 signaling is implicated in multiple neurodegenerative conditions [8], including AD [9], Parkinson’s disease [10], and multiple sclerosis [11].

The TNFR1 signaling pathway has been identified as a key regulator of both apoptosis and necroptosis. Binding of TNF to TNFR1 induces a conformational change in the receptor-ligand complex that enables the recruitment of TNFR1-associated death domain (TRADD) and receptor-interacting protein kinase 1 (RIPK1) and the formation of downstream complexes I and II [12,13]. While complex I triggers the nuclear factor-κB (NF-κB) pathway responsible for inducing cytokine production and anti-apoptotic gene expression, complex II mediates cell death pathways including apoptosis and necroptosis, which is dependent on caspase-8 activity [14]. Caspase-8 function is regulated by its phosphorylation state, expression level, and interaction with the autolysosomal network or the ubiquitin-proteosome system [15]. In situations where caspase-8 is absent or inactive, RIPK1 is not cleaved, leading to the phosphorylation of RIPK1, RIPK3, and mixed-lineage kinase domain-like protein (MLKL). This results in the formation of phosphorylated MLKL (p-MLKL) oligomers, initiating the process of necroptosis. Rupturing of the cell membrane by p-MLKL oligomers and the subsequent release of intracellular components associated with damage-associated molecular patterns (DAMPs) are characteristic hallmarks of necroptotic cell death [14] (**Fig. 1A**).

**Fig. 1.**
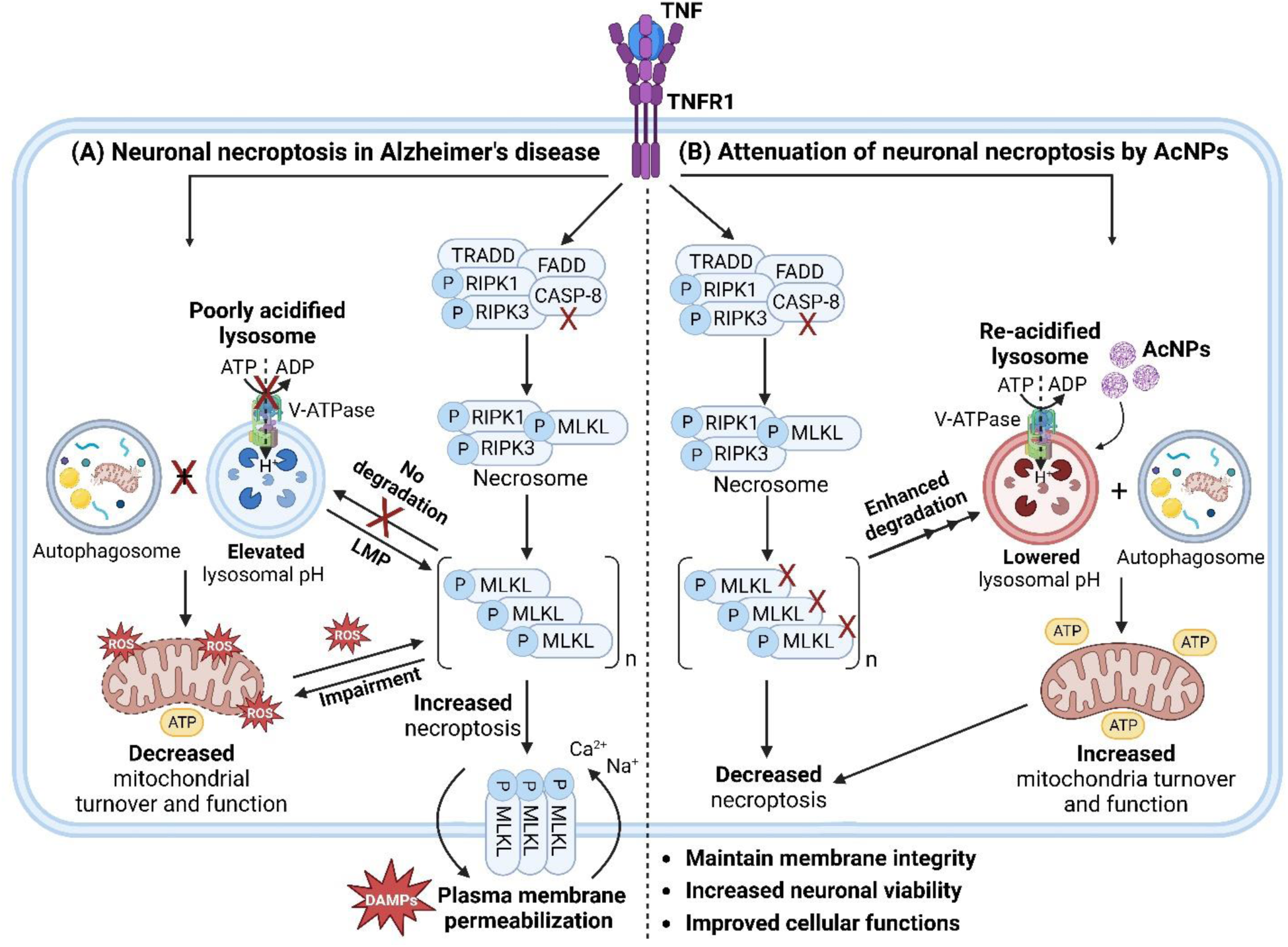
Schematic illustration of TNF-TNFR1 mediated lysosomal dysfunction and neuronal necroptosis. (A) Under TNF stimulation, necroptosis pathway is activated and autolysosomal function is impaired due to defective lysosomal acidification. This results in a reduction in autophagic degradation of necrosome molecules such as toxic p-MLKL oligomers and a decrease in turnover of damaged mitochondria under inflammatory condition. Lysosomal membrane permeabilization (LMP) and the generation of ROS by impaired mitochondria further propagate p-MLKL oligomerization. The accumulated p-MLKL oligomers rupture the plasma membrane, resulting in necroptotic neuronal death and leading to the release and exposure of damage-associated molecular patterns (DAMPs) which are regarded as endogenous danger signals. (B) Lysosome-acidifying nanoparticles (AcNPs) restore lysosomal acidification by lowering luminal pH and promote autophagic degradation under TNF stimulated condition. This leads to enhanced autophagic degradation of p-MLKL oligomers, increased turnover of damaged mitochondria, and improved mitochondrial functions, contributing to decreased necroptosis. Overall, restoration of lysosomal acidification under TNF stimulated inflammatory condition attenuates necroptosis and improves neuronal functions and viability. The figure was created with BioRender.com.

Mounting evidence has shown that both TNF and TNFR1 expression are increased in post-mortem human AD brains [16,17] and the levels of TNF and TNFR1 in the cerebrospinal fluid (CSF) and plasma are elevated [18,19]. Studies in post-mortem human AD brains have revealed the role of TNFR1 activation in inducing neuronal necroptosis as characterized by specific expression of p-MLKL in degenerating neurons without activation of apoptosis [17,20,21]. Specifically, our recent study has demonstrated that these degenerating neurons are expressing phosphorylated tau and located near Aβ plaques, activated microglia, and cytotoxic T-cells, indicating that the neuroinflammatory microenvironment renders neurons susceptible to death [17]. In transgenic mouse models of familial AD [20,22–24] and sporadic mouse model with Aβ oligomers injection [25], the elevated levels of necrosome molecules reinforce the mechanistic role of necroptosis in AD pathogenesis. Furthermore, stereotaxic injection of TNF in wild-type (WT) mice has been reported to activate necroptosis in mouse hippocampus [26]. In neuronal culture, studies have also shown that TNF activates necroptosis, with the highest percentage of cell death observed under the condition when caspases and inhibitor of apoptosis proteins (IAPs) are inactivated [17,24,26,27]. Lowering the level of TNFR1 and treatment of necroptosis inhibitors or anti-TNF antibodies are able to prevent necroptotic neuronal death, reduce the accumulation of toxic protein aggregates, and rescue cognitive impairments in neurons and AD mouse models [20,22–24,26,27]. In addition to the presence of necroptosis in AD, recent studies have also highlighted the role of lysosomal dysfunction and autophagic impairment in TNFR1 mediated neuronal death [28,29]. The autolysosome pathway initiates with phagophore and autophagosome formation followed by fusion of autophagosome and lysosome to form autolysosome which requires lysosomes with sufficient luminal acidification to have a complete autophagic degradation process [30]. It is important to note that while TNF has been suggested to promote autophagy initiation, studies have shown an increased accumulation of autophagic vesicles, indicating a terminal failure in the autophagic process [27,31]. This is supported by strong evidence of lysosomal acidification dysfunction [24,27,32] and lysosomal membrane permeabilization (LMP) [28] under TNF stimulated condition. While V-ATPase dysfunction is a key culprit contributing to defective lysosomal acidification [33], reactive oxygen species (ROS) associated with mitochondrial dysfunction also drives LMP and leads to cathepsin release and cleavage of caspases, exacerbating necroptosis and cellular disintegration [28,34]. Furthermore, interaction between TNFR1 signaling axis and autolysosomal pathway and the accumulation of necrosome molecules (e.g., p-MLKL) in poorly acidified lysosomes play key roles in mediating neuronal necroptosis [24,35]. Altogether, these studies suggest that restoration of lysosomal acidification and function is a viable strategy to inhibit TNFR1 mediated neuronal necroptosis (**Fig. 1B**).

In this study, we revealed using neuropathological analysis of human post-mortem AD brains that TNFR1 mediated necroptosis correlates with autolysosomal dysfunction as indicated by downregulation of V-ATPase subunits and accumulation of autophagy receptor p62. Specifically, we showed that TNF induces lysosomal acidification defect, disruption of lysosomal membrane dynamics, LMP, autophagic impairment, and neuronal death in SH-SY5Y neuroblastoma cells. Importantly, we further demonstrated that lysosome-acidifying nanoparticles (AcNPs) restore lysosomal acidification and membrane homeostasis, improve autophagic and mitochondrial function, and rescue neuronal necroptosis in both TNF-treated SH-SY5Y cells and APP^NL-G-F^ mouse model of AD. We propose that targeting lysosomal acidification dysfunction represents a novel therapeutic strategy to attenuate neuronal necroptosis in AD.

## Materials and Methods

### Data mining of human AD transcriptomic dataset

The RNA-sequencing (RNA-seq) transcriptomic dataset GSE173955 [36] was obtained from the Gene Expression Omnibus (GEO) database [37]. The 10 healthy control (HC) and 8 AD samples are from the hippocampus region of post-mortem brains with the age of the HC samples at 77.00 ± 8.50 years and AD samples at 91.75 ± 6.08 years. The dataset was analyzed using the GEO2R tool that makes use of the limma package in the Rstudio [38]. The mRNA expression from AD samples were compared against HC samples, and the differentially expressed genes (DEGs) were identified using the parameters of false discovery rate (FDR) adjusted *P*<0.01 and log_2_(fold change (FC))≥|0.5|. The pathway enrichment analysis was conducted using the Database for Annotation, Visualization and Integrated Discovery (DAVID) [39] which provides functional annotations of the DEGs based on Kyoto Encyclopedia of Genes and Genomes (KEGG), Gene Ontology Biological Process (GOBP), and Gene Ontology Cellular Component (GOCC) databases. In addition, gene set enrichment analysis (GSEA) [40] was conducted for both AD and HC samples of the GSE173955 dataset by comparing to a list of gene sets following default settings including 1000 permutations, Signal2Noise metric for ranking genes, and weighted enrichment statistics. The chip platform “Human_Gene_Symbol_with_Remapping_MSigDB.v2023.1.Hs.chip” was used for gene identification. The GOBP gene set from the Molecular Signatures Database (MSigDB) [41] was used as a reference.

### Human post-mortem brain specimens

The human post-mortem AD brains used in this study were sourced from the Multiple Sclerosis and Parkinson’s Tissue Bank at Imperial College London and the South West Dementia Brain Bank at the University of Bristol. Snap-frozen tissue blocks from the hippocampus were acquired from 10 AD cases (82.90 ± 7.88 years) and 10 HC cases (84.20 ± 7.39 years). The post-mortem donations were obtained with full informed consent and under ethical approval by the National Research Ethics Committee (08/MRE09/31 and NHS REC No 18/SW/0029). The use of human samples in this study is also approved by the Institutional Review Board (IRB-2023-143) from the Nanyang Technological University (NTU), Singapore. Demographic data and neuropathological characteristics of the brain samples (**Table S1**) were provided by both brain banks, following standardized criteria of the UK MRC Brain Bank Network.

### Animal maintenance and stereotaxic intrahippocampal injection

We used WT and APP^NL-G-F^ homozygous knock-in mice (both male and female) with C57BL/6 background at 3-month-old for treatment procedures and the mice were sacrificed at 4-month-old for experiments. WT and APP^NL-G-F^ mice were bred at the animal facility. The mice were housed in a temperature-controlled room (25 °C) in virus-free facilities on a 12 h light/dark cycle (7:30 a.m. on/7:30 p.m. off) and water ad libitum. All mice were maintained under specific pathogen-free conditions and treated with humane care. In each animal experiment, mice were randomly allocated to receive either PBS or AcNPs (75 mg/mL) at a total volume of 2 µL (total amount of 150 µg) via stereotaxic intrahippocampal injection at the respective coordinates from bregma (A/P: −2.1 mm; M/L: +/− 1.7 mm; D/V: −2.5 mm) [42]. For all stereotaxic injections, a 26G Hamilton syringe was used and the solution was applied at a speed of 0.5 µL/min. After injection, the needle was kept in place for an additional 5 min before gentle withdrawal. The mice were sacrificed 1 month post injection. Mice were sacrificed by perfusion and mouse brains were snap frozen in liquid nitrogen upon extraction and stored in −80°C for further processing. All procedures were conducted with the approval of the Institutional Animal Care and Use Committee (IACUC) animal use protocol A21043 and A22058 from NTU Singapore.

### Tissue fixation and sectioning

For animal experiments, mouse brains were first fixed in 4% paraformaldehyde (PFA) in PBS solution (VWR, Cat# ALFAJ61899.AK) for 24 h, followed by overnight fixation in 30% sucrose solution (Sigma-Aldrich, Cat# S0389). The samples were then embedded in Tissue-Tek O.C.T compound (Sakura Finetek USA, Cat# 4583) using molds measuring 15x15 mm (Epredia, Cat# 58950). The mouse brains were then cut into 20 µm thick frozen tissue sections using a cryostat (Leica, Cat# CM1950) and placed on Superfrost Plus Adhesion Microscopic glass slides (Epredia, Cat# J1800AMNZ). For use of the human post-mortem brain tissues, 20 µm thick cryosections were cut from the snap frozen hippocampal blocks and placed on Superfrost Plus Adhesion Microscopic glass slides.

### Tissue immunostaining and image acquisition

For immunostaining of both mouse and human brain tissues, the frozen sections were blocked with 5% normal goat serum (NGS) (Thermo Fisher Scientific, Cat# 31872) for 1 h at room temperature. Subsequently, the sections were incubated with primary antibodies followed by fluorescent secondary antibodies (**Table S2**). The tissue sections were subsequently incubated with DAPI (Thermo Fisher Scientific, Cat# 9542) for 10 min before washing with PBS. Lastly, antifade mounting medium (VectaShield, Cat# H-1000) was applied to the sections, before mounting of coverslip (24x50mm, Paul Marienfeld, Cat# 01-022-22). Slide scanner (Carl Zeiss Axio Scan Z1) was used to acquire fluorescent images at 40X magnification. Zen Blue software v2.6 was used to analyze the acquired images.

### AcNPs synthesis and characterizations

The AcNPs were synthesized from poly(ethylene tetrafluorosuccinate-co-succinate) PEFSU polyesters as described previously [43]. Briefly, the polyesters (25% PEFSU) were dissolved in acetonitrile (Sigma Aldrich, Cat # 271004) and then passed through a 0.2 µm syringe filter (Millipore, Cat# Z741696) to eliminate any particles. Sodium dodecyl sulfate (SDS) (Sigma Aldrich, Cat# L3771) was dissolved in MilliQ water and stirred vigorously at 1700 rpm. The acetonitrile polyester solution was slowly added drop by drop into the vigorously stirred aqueous SDS solution. Once all the polyester solution had been added, the resulting emulsion was transferred into SnakeSkin dialysis tubing (MWCO 10 kDa) (Thermofisher Scientific, Cat# 68100) and dialyzed against MilliQ water for 24 h. For dynamic light scattering (DLS) measurements, 200 µL of the solution was diluted in 2.8 mL of MilliQ water, and the diameter was measured using the Brookhaven 90 plus NP sizer DLS instrument. For scanning electron microscopy (SEM) characterization, the AcNPs were diluted 50-fold in MilliQ water, and 10 µL droplets were applied onto silicon wafers, allowing them to air dry overnight. Later, these wafers were affixed to aluminium stubs using copper tape and coated with a 5 nm-thick Au/Pd layer through sputter coating. The samples were analyzed using a Supra 55VP field emission scanning electron microscope (ZEISS) with an accelerating voltage of 2 kV and a working distance of 5.5 cm.

### Cell culture and cytotoxicity assay

SH-SY5Y cells (ATCC, Cat# CRL-2266) were cultured in DMEM/F12 media (Thermofisher scientific, Cat# 11320033) supplemented with 10% fetal bovine serum (Gibco FBS HI, Cat# A5256701) and 1% penicillin-streptomycin (Thermofisher Scientific, Cat# 15140122). The cytotoxicity assay was evaluated using an MTS Assay Kit (Abcam, Cat# ab197010). SH-SY5Y cells were cultured in a 96-well plate at 15000 cells/well for 24 h and subsequently treated with and without TNF and the respective doses of the AcNPs for 24 h. The cell viability was quantified relative to the no treatment control.

### Lysosomal pH measurements

For live-cell imaging of lysosomal pH, SH-SY5Y cells subjected to respective treatments were stained with LysoSensor™ Yellow/Blue DND-160 (Thermofisher Scientific, Cat# L7545) following manufacturer’s protocol for 15 min. The stained cells were then imaged using a confocal microscope (ZEISS) with 360 nm excitation and images were acquired at 440 nm (blue channel) and 540 nm (yellow channel). The ratio between yellow and blue fluorescence intensities was calculated, and pH quantification was achieved by comparing it to a standard curve with known pH values. The standard curve was generated by correlating LysoSensor fluorescence ratio to known pH values using 2-(N-morpholino) ethanesulfonic acid buffer with varying pH.

### Cathepsins activity assay

For cathepsin B activity assay, SH-SY5Y cells subjected to respective treatments were incubated with Magic red cathepsin B (MR-cathepsin B, Immunochemistry Technologies, Cat# 937) following manufacturer’s protocol for 1 h. For cathepsin D activity assay, the SH-SY5Y cells were treated according to manufacturer’s instructions (Abcam, Cat# ab65302). After treatment, the cells were washed three times with PBS, and the fluorescence was measured using a Synergy H1 model of hybrid multi-mode plate-reader with excitation/emission wavelengths of 531/629 nm for cathepsin B activity and 328/460 nm for cathepsin D activity.

### Cell immunofluorescence and image acquisition

For cell immunofluorescence imaging, SH-SY5Y cells were first subjected to respective treatment conditions. Subsequently, the cells were fixed using a 4% paraformaldehyde (PFA) in PBS solution for 20 min and rinsed twice with PBS. The fixed cells were washed three times with 0.1% Triton X-100 (Sigma Aldrich, Cat# X100) in PBS. Next, the cells were blocked with 5% normal goat serum (NGS) (Thermofisher Scientific, Cat# 31872) for 1 h at room temperature. The cells were then incubated with primary antibodies listed under **Table S2** overnight at 4 °C. The next day, cells were washed three times with 0.1% Triton X-100 in PBS, the cells were treated with fluorescent secondary antibodies listed under **Table S2** for 1 h at room temperature. Subsequently, the cells were washed with PBS. DAPI staining (Thermofisher Scientific, Cat# 9542) was performed for 10 min, followed by PBS washes. Fluorescent images were captured using a confocal microscope (ZEISS) at blue (excitation/emission 405/460 nm), green (excitation/emission 490/520) and red (excitation/emission 561/617 nm) channels.

### Autophagy and mitochondrial turnover reporter assays

SH-SY5Y cells were transfected with GFP-LC3 reporter plasmid (Addgene, Cat# 11546) for 24 h, before treatment with different conditions. The cells expressing GFP-LC3 were then imaged using confocal microscopy (excitation/emission 490/510 nm) to visualize GFP-LC3 punctate dots within cells. To quantify mitochondria turnover under different conditions, SH-SY5Y cells were first transfected with mCherry-GFP-FIS1 (University of Dundee, Cat# DU55501) for 24 h to express the mitophagy reporter. Subsequently, the cells were subjected to different treatment conditions for another 24 h. Before imaging the cells, the LysoSensor™ Blue DND-167 dye (Thermofisher Scientific, Cat# L7533) was added for 15 min. Confocal microscopy was used to capture fluorescent images at blue (excitation/emission 405/460 nm), green (excitation/emission 490/520 nm), and red (excitation/emission 590/610 nm) channels.

### Characterization of mitochondrial morphology and functions

SH-SY5Y cells were subjected to respective treatment conditions and then stained with MitoTracker Deep Red (Thermofisher Scientific, Cat# M22426) following manufacturer’s protocol for 30 min. The stained cells were washed and subsequently imaged using confocal microscopy with excitation/emission wavelengths of 640/665 nm. For mitochondrial morphology analysis, the acquired confocal images of mitochondria were processed using the Mitochondrial Network Analysis (MiNA) ImageJ macro [44]. Quantifiable measurements of mitochondrial morphology, such as mitochondrial footprint (µm^2^) and network branch length (µm), were plotted.

The processing parameters were kept consistent across images acquired from different treatment conditions. To measure mitochondrial membrane potential (MMP), SH-SY5Y cells were cultured in a 96-well plate and subjected to respective treatments, followed by analysis using the TMRE-Mitochondrial Membrane Potential Assay Kit (Abcam, Cat# ab113852). To measure mitochondrial superoxide release, the Mitochondrial Superoxide Assay Kit (Abcam, Cat# ab219943) was used. Both assays were conducted following the protocols provided by the manufacturer. The measurements were obtained using a Synergy H1 plate-reader with excitation/emission wavelengths of 549/575 nm for TMRE and 510/610 nm for the superoxide signals.

### Western blotting

Sample preparations for cell lysates as well as mouse and human brain lysates were performed with the following procedures. SH-SY5Y cells treated with respective conditions were first washed with PBS, followed by the addition of lysis solution consisting of RIPA buffer (Thermofisher Scientific, Cat# 89900) supplemented with protease and phosphatase inhibitors (Thermofisher Scientific, Cat# 78440). Mouse whole brain tissues were homogenized using similar lysis solution as described above. The human hippocampus brain tissues were carefully extracted from frozen human HC and AD brain samples by delicately scoring the tissue block and then isolating the tissue through cryosectioning in a Leica cryostat. The resulting tissue samples were then homogenized using the lysis solution. All lysates were kept on ice for 15-30 min followed by centrifugation for 10 min at 13,500 g at 4 °C.

Once the cell and tissue lysates were prepared, Pierce™ BCA Protein Assay Kit (Thermofisher Scientific, Cat# 23225) was used to determine total protein concentrations for each sample. Samples were then mixed with 4X Laemmli Sample Buffer (Bio-Rad, Cat# 1610747) and boiled at 95 °C for 5 min before loading and running in 4-15% Mini-PROTEAN TGX Precast Protein Gels (Bio-Rad, Cat# 4561083) with Precision Plus Protein Dual Color Standards (Bio-Rad, Cat# 1610374). The proteins were transferred to polyvinylidene fluoride (PVDF) membrane (Millipore, Cat# ISEQ00010), and blocked with blocking buffer (Bio-Rad, Cat# 1706404). The membranes were then incubated with respective primary and secondary antibodies (**Table S2**). Densitometry was performed using ImageJ and protein expression levels were normalized to GAPDH loading control.

### Statistical analysis

Statistical analyses were performed using the GraphPad Prism 11 software. Statistical analysis was conducted by using unpaired Student’s t test for difference between two conditions and one-way ANOVA with post hoc Tukey’s test for multiple comparisons. Statistical significance was determined by *P*<0.05 and indicated by \**P*<0.05, \*\**P*<0.01, \*\*\**P*<0.001, \*\*\*\**P*<0.0001 or ns for non-significance.

## Results

### Correlation of TNFR1 mediated necroptosis and autolysosomal dysfunction in hippocampal region of human AD brains

To investigate whether defective lysosomal acidification is present in human AD brains, we conducted data mining to analyze an existing RNA-seq transcriptomic dataset (GSE173955) [36] containing mRNA expression of hippocampus tissues from 8 AD and 10 HC samples. With a cutoff of FDR adjusted *P*<0.01 and log_2_(fold change (FC))>|0.5|, we identified 1769 DEGs with 1070 downregulated genes and 699 upregulated genes (**Fig. 2A**). We then performed pathway enrichment analysis using KEGG, GOBP, and GOCC databases to obtain the functional annotations of the DEGs. First, the identified DEGs were associated with pathways of neurodegeneration, AD, as well as glutamatergic and GABAergic synapses, validating the usefulness of this dataset for our study (**Fig. 2B**). We also showed that pathways related to synaptic functions (e.g., synaptic vesicle cycle and synaptic transmission) and metabolic activities (e.g., ROS and oxidative phosphorylation) are particularly enriched (**Fig. 2B-D**). Furthermore, our analysis highlighted the association of DEGs to lysosomal membrane, V-ATPase, proton transmembrane transport, vacuolar acidification, and regulation of TRP channels (**Fig. 2B-D**).

**Fig. 2.**
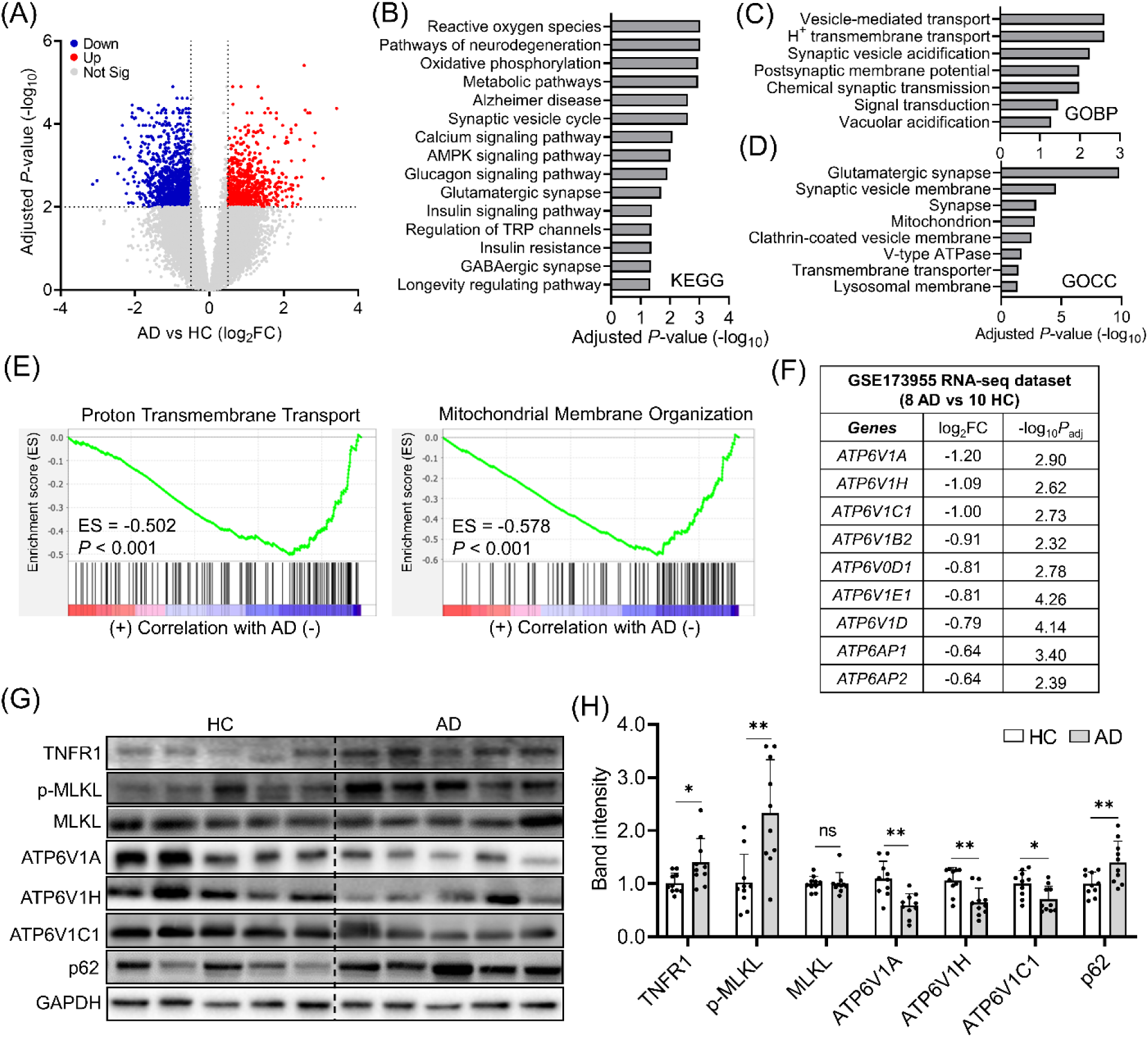
Downregulation of V-ATPase subunits required for lysosomal acidification in human AD brains. (A) Volcano plot illustrating the downregulated (blue) and upregulated (red) DEGs from the GSE173955 RNA-seq transcriptomic dataset. The analysis was conducted based on the mRNA expression of the 8 AD and 10 HC brain tissues using a cutoff of FDR adjusted *P*<0.01 and log_2_FC>|0.5|. (B-D) Pathway enrichment analyses performed using the (B) KEGG, (C) GOBP, and (D) GOCC databases, illustrating the functional annotations of the DEGs. (E) GSEA analysis of the GSE173955 dataset showing downregulation of proton transmembrane transport and mitochondrial membrane organization in AD compared to HC brains. (F) Specific V-ATPase subunits that are identified to be downregulated in AD brains. (G-H) Western blotting analysis of TNFR1 mediated necroptosis (TNFR1, p-MLKL, MLKL), top three downregulated V-ATPase subunits (ATP6V1A, ATP6V1H, and ATP6V1C1), and autophagy receptor p62 in the human HC and AD brains with GAPDH as a loading control. Data presented are relative to HC. Data are means ± SD of N=10 HC and N=10 AD brains. \**P* < 0.05, \*\**P* < 0.01, and ns indicates non-significance by unpaired Student’s t test for comparison between two samples.

To understand how the genes are involved in these pathways, we performed the pathway enrichment analysis separately for the upregulated and downregulated DEGs. From the analysis, upregulated genes were mainly related to regulation of RNA transcription. The analysis for downregulated genes suggests impairments in synaptic and metabolic activities as well as mitochondrial, autophagic, and lysosomal dysfunctions in AD (**Fig. S1A-C**). We then conducted GSEA analysis and illustrated downregulation of proton transmembrane transport, mitochondrial activities, and synaptic functions (**Fig. 2E and S1D**). With a focus in understanding lysosomal acidification alterations, we identified nine lysosomal V-ATPase subunits (*ATP6V1A*, *ATP6V1H*, *ATP6V1C1*, *ATP6V1B2*, *ATP6V0D1*, *ATP6V1E1*, *ATP6V1D*, *ATP6AP1*, and *ATP6AP2*) to be significantly downregulated in the dataset (**Fig. 2F**), indicating that lysosomal acidification was impaired in AD brains.

We further confirmed these observations in the protein level using 10 HC and 10 AD post-mortem human brains (**Table S1**). As an initial control, we illustrated the presence of activated microglia (**Fig. S2A**) as well as increased TNF and phospho-tau levels (**Fig. S2B-C**) in AD brains as compared to controls, indicative of neurodegenerative pathology. We then showed that there is an increase in both TNFR1 and p-MLKL levels with no significant changes in total MLKL level (**Fig. 2G-H**) and caspase activities (**Fig. S2B-C**), confirming the presence of TNFR1 induced necroptosis in AD brains. We further demonstrated reduction in the protein levels of the top three downregulated (log_2_FC≤-1) V-ATPase subunits (ATP6V1A, ATP6V1H, and ATP6V1C1) that we have identified from data mining, together with an increased accumulation of p62, illustrating both lysosomal dysfunction and autophagic impairment in AD brains (**Fig. 2G-H**). We further observed the expression of TNFR1, p-MLKL, MLKL, V-ATPase subunits and p62 in hippocampal neurons of human HC and AD brains (**Fig. S3**), thus establishing the correlation between TNFR1 induced neuronal necroptosis and autolysosomal dysfunction.

### TNF induces lysosomal pH elevation in SH-SY5Y cells and AcNPs restore lysosomal acidification and proteolytic function

To investigate the role of TNF-TNFR1 signaling axis in autolysosomal function, we first determined the effect of TNF in altering lysosomal acidification in SH-SY5Y cells. We showed that V-ATPase subunits (ATP6V1A, ATP6V1H, and ATP6V1C1) are decreased in TNF treated cells (**Fig. 3A-B**), recapitulating their downregulation in human AD brains. As expected, TNF induces lysosomal pH elevation from 4.5 to 5.6 in SH-SY5Y cells, making this cell model suitable to study lysosomal acidification dysfunction (**Fig. 3C-D**). Importantly, this lysosomal pH elevation is specific to TNF induction as stimulation by other cytokines did not change lysosomal pH (**Fig. 3C-D**). We then utilized a new type of lysosome-acidifying nanoparticles (AcNPs) that are capable of efficiently localizing into impaired lysosomes (**Fig. S4A**) [43,45], to directly restore organelle luminal acidification in TNF stimulated SH-SY5Y cells. In control SH-SY5Y cells, rhodamine-labeled AcNPs (Rho-AcNPs) localized to Lysosensor-positive lysosomes (**Fig. 3E**). AcNPs did not alter lysosomal pH in control cells with normally acidified lysosomes or induce detectable toxicity (**Fig. S4B-C**). Importantly, we demonstrated that AcNPs acidify impaired lysosomes in a dose dependent manner (NP50 = 50 µg/mL and NP100 = 100 µg/mL) by lowering their pH from 5.6 to 4.6 (**Fig. 3F-G**). Consistent with the importance of acidification for lysosomal proteolytic function, TNF also reduced cathepsin B and D activities in SH-SY5Y cells (**Fig. 3H-I**) whereas AcNP treatment restored both enzyme activities.

**Fig. 3.**
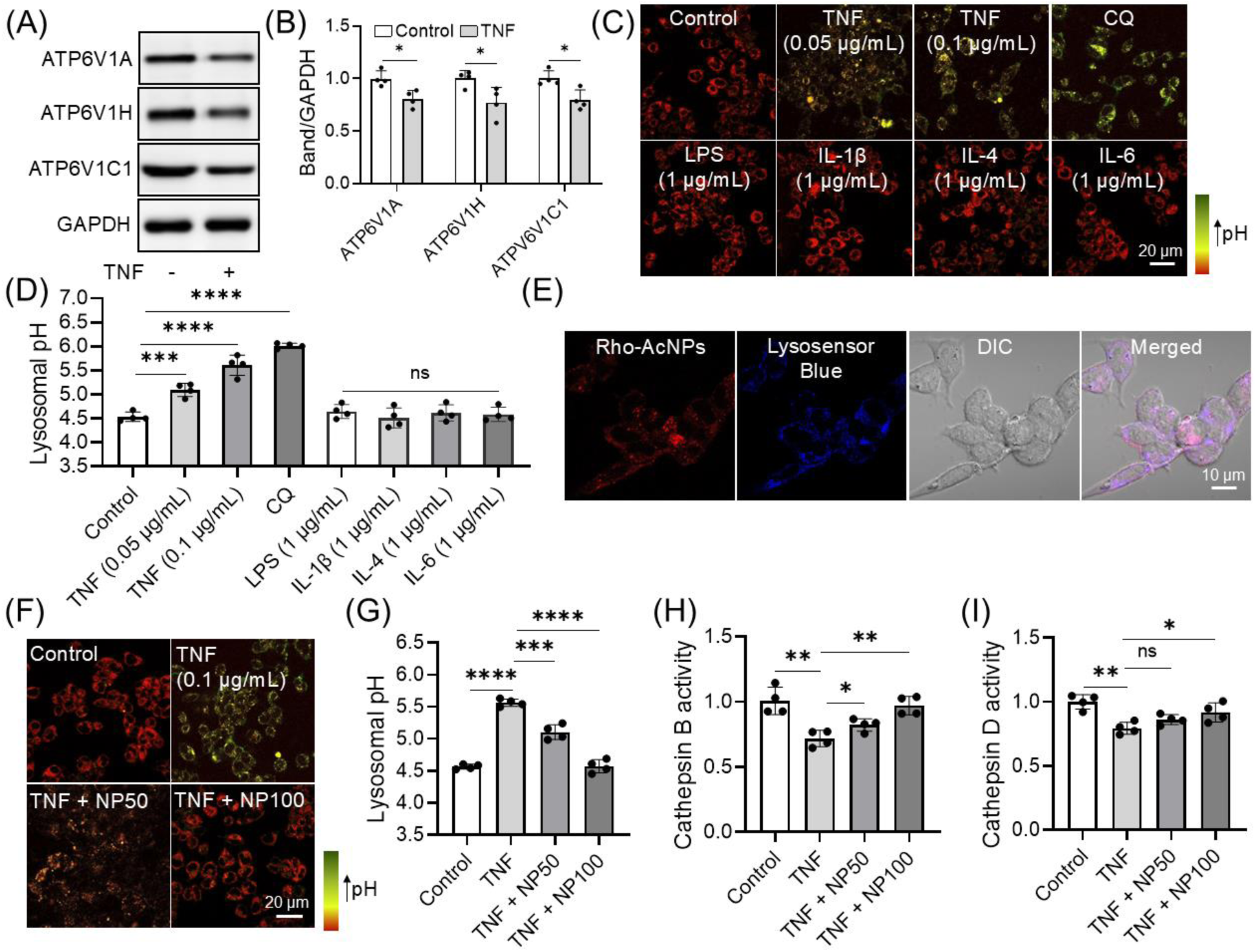
Reduced expression of V-ATPase subunits and lysosomal acidification dysfunction in SH-SY5Y cells under TNF stimulated condition. (A-B) Western blotting analysis and quantification of V-ATPase subunits (ATP6V1A, ATP6V1H, and ATP6V1C1) in TNF-treated SH-SY5Y cells with GAPDH as a loading control. (C-D) Lysosomal pH measurement and quantification using LysoSensor Yellow/Blue in SH-SY5Y cells following stimulation with TNF or other cytokines. (E) Representative confocal images showing localization of rhodamine-labeled AcNPs (Rho-AcNPs) to Lysosensor-positive lysosomes in control SH-SY5Y cells. (F-G) Lysosomal pH measurement and quantification in TNF-treated SH-SY5Y cells following treatment with AcNPs at 50 μg/mL (NP50) or 100 μg/mL (NP100). (H-I) Lysosomal cathepsin B and D activities in SH-SY5Y cells under the indicated treatment conditions. Data are presented relative to untreated control cells without TNF or AcNPs treatment. Data are means ± SD of N = 4 independent experiments. *P* < 0.05, \**P* < 0.01, \*\**P* < 0.001, \*\*\**P* < 0.0001, and ns, not significant, by unpaired Student’s *t*-test for comparisons between two groups and one-way ANOVA with Tukey’s post hoc test for multiple comparisons.

### TNF induces lysosomal membrane damage and disrupts lysosomal membrane homeostasis

TNF induced LMP as indicated by increased galectin-3 puncta per cell (**Fig. 4A**). Because lysosomal membrane injury activates coordinated repair and quality-control responses, we next examined components of these pathways. ESCRT-dependent membrane repair represents an early response to lysosomal membrane damage, whereas persistent damage can promote lysophagic clearance of irreparably damaged lysosomes [46,47]. TNF increased CHMP2B, an ESCRT-associated membrane repair protein, consistent with activation of a lysosomal membrane damage response (**Fig. 4B**) [17]. TNF also increased TRIM16, which participates in the recognition and autophagic clearance of damaged lysosomes (**Fig. 4C**) [48,49]. AcNPs treatment reduced both CHMP2B and TRIM16 toward control levels, corroborating with reduced LMP and restoration of lysosomal function (**Fig. 4B-C**), suggesting that re-acidification attenuates the lysosomal stress that activates these damage-response pathways. In contrast, TNF reduced LAPTM4B (**Fig. 4D**), a lysosomal membrane protein implicated in maintaining lysosomal membrane stability, acidification, and autophagic function [50,51]. AcNPs treatment restored LAPTM4B expression concomitant with reduced LMP (**Fig. 4A,D**). We next examined Arl8B (**Fig. 4E**) and VAMP7 (**Fig. 4F**), which regulate complementary aspects of lysosomal membrane dynamics, including lysosomal positioning and trafficking, and membrane fusion, respectively [52–54]. TNF reduced expression levels of both proteins, whereas AcNPs treatment restored their expression (**Fig. 4E-F**), suggesting that inflammatory stress also disrupts lysosomal trafficking and membrane dynamics. These findings demonstrate that TNF induces coordinated lysosomal membrane damage and disruption of membrane homeostasis, accompanied by activation of lysosomal damage-response pathways and altered trafficking and fusion machinery. AcNPs-mediated re-acidification attenuates these defects and restores lysosomal membrane homeostasis.

**Fig. 4.**
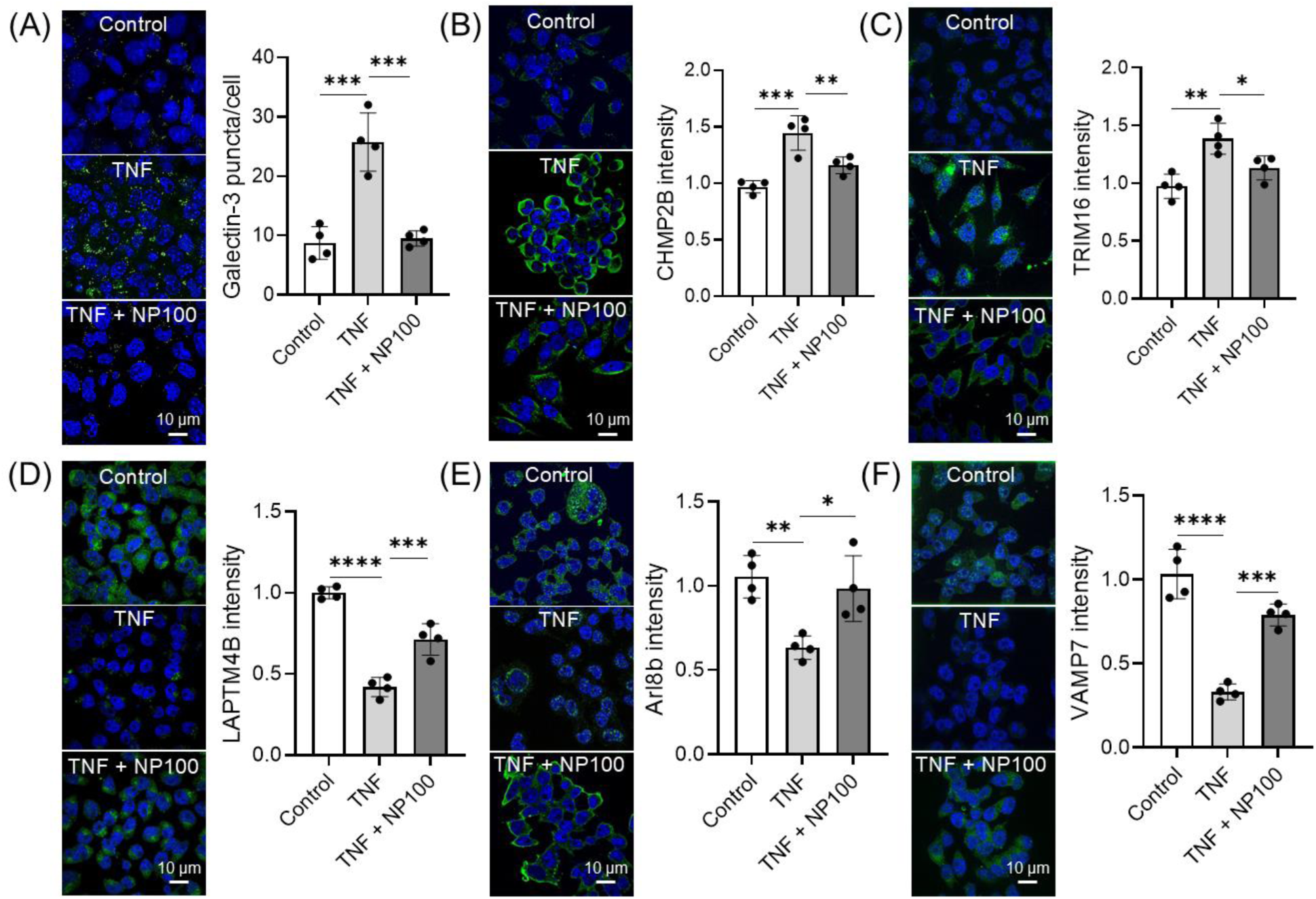
AcNPs attenuate TNF-induced lysosomal membrane damage and restore lysosomal membrane dynamics. (A) Representative images and quantification of galectin-3 puncta as a marker of LMP in TNF-treated SH-SY5Y cells. (B-C) Representative images and quantification of the lysosomal damage-response proteins CHMP2B and TRIM16, respectively, following TNF stimulation and AcNPs treatment. (D) Representative images and quantification of LAPTM4B, a lysosomal membrane protein involved in lysosomal membrane homeostasis. (E-F) Representative images and quantification of Arl8B and VAMP7, regulators of lysosomal positioning and trafficking and lysosomal membrane fusion, respectively. Data are presented relative to control cells without TNF or AcNP treatment. NP100 indicates AcNPs treatment at 100 µg/mL. TNF was used at 0.1 µg/mL unless otherwise stated. Data are means ± SD of N=4 independent experiments. \**P* < 0.05, \*\**P* < 0.01, \*\*\**P* < 0.001, and \*\*\*\**P* < 0.0001 by one-way ANOVA with post hoc Tukey’s test for multiple comparisons.

### AcNPs promote autophagic activity and increase mitochondrial turnover in TNF stimulated SH-SY5Y cells

As lysosome degradation is the final and determining step of complete autophagic degradation, we examined whether re-acidification of impaired lysosomes by AcNPs in TNF stimulated SH-SY5Y cells restores autophagic function. TNF caused autophagy inhibition as illustrated by the accumulation of autophagy-associated proteins p62 and LC3II in SH-SY5Y cells, and AcNPs treatment promoted their clearance (**Fig. 5A-C**). We further confirmed this observation by using a GFP-LC3 reporter, where TNF induced an aggregation of LC3 puncta and treatment of AcNPs increased the degradation and clearance of these puncta (**Fig. 5D-E**). Functional autophagy is required for proper turnover of mitochondria and maintenance of normal mitochondrial functions. To determine whether improved autophagic function with AcNPs treatment promotes mitochondrial turnover in TNF-treated SH-SY5Y cells, we transfected the cells with a mCherry-GFP-FIS1 reporter plasmid that reflects the accumulation of mitochondria in lysosomes with different extent of acidification by quantifying the fluorescence signals [55]. The green fluorescent protein (GFP) signal is quenched when mitochondria colocalize with sufficiently acidified lysosomes with low pH but the green fluorescence remained high in impaired lysosomes with elevated pH. Under all conditions, the mCherry fluorescence signal is unchanged and acts a control for the presence of mitochondria. In untreated SH-SY5Y control cells, normal mitochondrial turnover was illustrated by a basal red and green fluorescence of the mCherry-GFP-FIS1 reporter. TNF treatment increased the green fluorescence intensity (**Fig. 5F-G**), indicating the accumulation of mitochondria in poorly acidified lysosomes. Their colocalization was indicated by white puncta formed by overlapping red (mitochondrial protein), green (elevated lysosomal pH), and blue (lysosomes) fluorescence. Treatment of AcNPs restores lysosomal acidification and quenches the GFP fluorescence (**Fig. 5F-G**), leading to the appearance of purple puncta when mitochondria (red) colocalize with lysosomes (blue). This result indicates that AcNPs treatment improves autophagic clearance and mitochondrial turnover in TNF-treated SH-SY5Y cells.

**Fig. 5.**
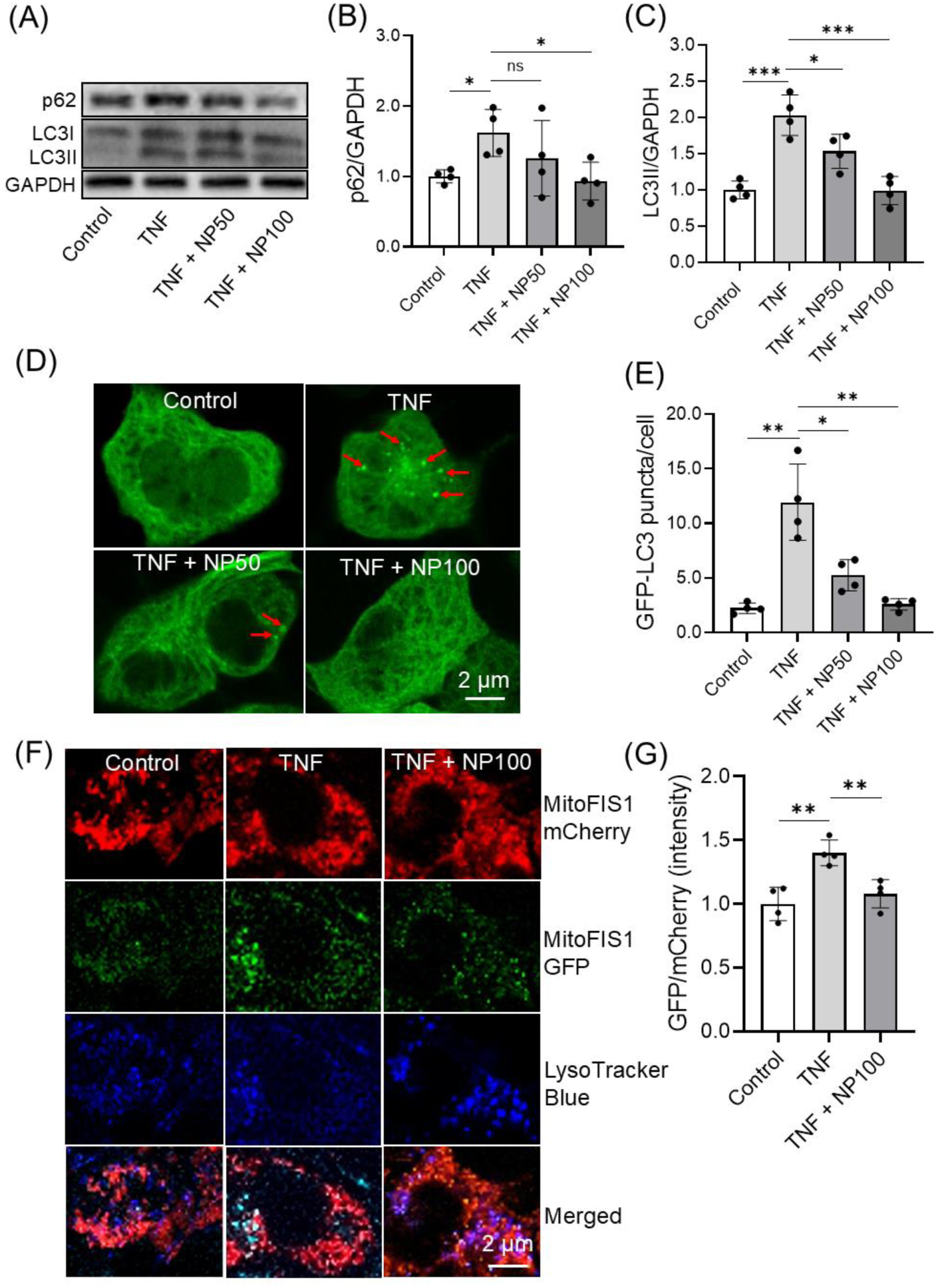
Re-acidification of lysosomes promotes autophagic degradation and mitochondrial turnover in TNF-treated SH-SY5Y cells. (A-C) Western blotting analysis and quantification of autophagy-associated proteins p62 and LC3II in SH-SY5Y cells under respective treatment conditions with GAPDH as a loading control. (D-E) Confocal microscopy images and quantification of GFP-LC3 reporter assay in SH-SY5Y cells under respective treatment conditions. (F-G) Confocal microscopy images and quantification of SH-SY5Y cells transfected with mCherry-GFP-FIS1 mitophagy reporter plasmid followed by respective treatments. Data presented are relative to control without TNF and AcNPs treatment. NP50 and NP100 indicate AcNPs treatments at 50 µg/mL and 100 µg/mL respectively. TNF was used at 0.1 µg/mL unless otherwise stated. Data are means ± SD of N=4 independent experiments. \**P* < 0.05, \*\**P* < 0.01, \*\*\**P* < 0.001, and ns indicates non-significance by one-way ANOVA with post hoc Tukey’s test for multiple comparisons.

### AcNPs improve mitochondrial function and rescue TNF-induced necroptosis in SH-SY5Y cells

With an increase in mitochondrial turnover, we then examined whether AcNPs restore mitochondrial functions by determining changes in their morphology, MMP, and ROS production. By staining with MitoTracker Deep Red, TNF induced more mitochondrial fragmentation (**Fig. 6A**) in SH-SY5Y cells as quantified by the decrease in mitochondrial footprint (**Fig. 6B**) and network branches (**Fig. 6C**), which are associated with mitochondrial dysfunctions [56,57]. With AcNPs treatment, there were more elongated mitochondria with tubular network (**Fig. 6A-C**), indicating an improvement in mitochondrial functions. Furthermore, AcNPs restored MMP as illustrated by the increase in TMRE intensity (**Fig. 6D**) and reduced ROS generation (**Fig. 6E**) in TNF-treated SH-SY5Y cells. Finally, we investigated whether AcNPs rescue TNF induced cell death in SH-SY5Y cells. TNF caused a significant cell death leading to 60% of cell viability (**Fig. 6F**) and treatment of AcNPs rescues cell death in a dose-dependent manner with a half maximal effective concentration (EC_50_) of 46 µg/mL, resulting in an increase in cell viability to 90% (**Fig. 6G**). As a control for specificity, we showed that AcNPs do not rescue lipopolysaccharides (LPS) induced cell death in SH-SY5Y cells (**Fig. S5A-B**) as the cellular impairment does not arise from lysosomal acidification dysfunction (**Fig. 3C-D**).

**Fig. 6.**
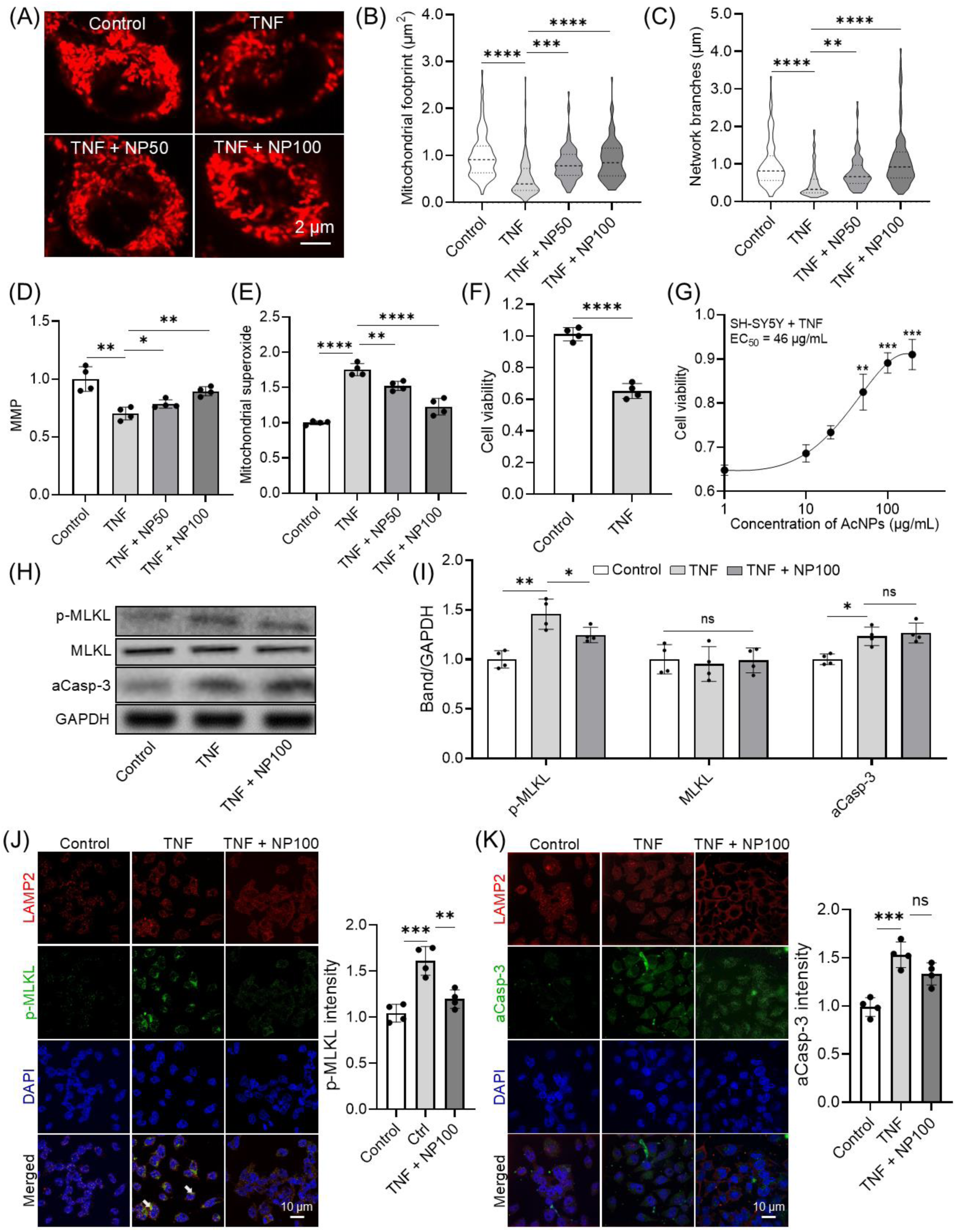
Restoration of lysosomal acidification improves mitochondrial function and rescues TNF induced neuronal necroptosis. (A) Confocal microscopy images of SH-SY5Y cells treated with respective conditions and stained with MitoTracker Deep Red. (B-C) Quantification of mitochondrial images using MiNA analysis to obtain measurements for (B) mitochondrial footprint and (C) network branches. (D-E) Characterizations of mitochondrial functions including the measurements of (D) mitochondrial membrane potential (MMP) and (E) reactive oxygen species (ROS) generation in SH-SY5Y cells under respective treatment conditions. (F) Measurement of cell viability of SH-SY5Y cells with and without TNF (100 ng/mL) treatment. (G) Measurement of cell viability of TNF-treated SH-SY5Y cells with addition of increasing doses of AcNPs (1 to 200 µg/mL). AcNPs rescue TNF induced cell death in a dose-dependent manner with an EC_50_ of 46 µg/mL. (H-I) Western blotting analysis and quantification of necroptosis markers (p-MLKL and MLKL) and caspase-3 activity (aCasp-3) in SH-SY5Y cells under respective treatment conditions with GAPDH as a loading control. (J) Representative immunofluorescence images and quantification of p-MLKL and (K) cleaved caspase-3 (aCasp-3) in TNF-treated SH-SY5Y cells with or without AcNP treatment. White arrows indicate co-localization of p-MLKL puncta with LAMP2 positive lysosomes. Data presented are relative to control without TNF and AcNPs treatment. NP50 and NP100 indicate AcNPs treatments at 50 µg/mL and 100 µg/mL respectively. TNF was used at 0.1 µg/mL unless otherwise stated. Data are means ± SD of N=4 independent experiments. \**P* < 0.05, \*\**P* < 0.01, \*\*\**P* < 0.001, \*\*\*\**P* < 0.0001, and ns indicates non-significance by unpaired Student’s t test for comparison between two samples and one-way ANOVA with post hoc Tukey’s test for multiple comparisons.

In TNF-treated SH-SY5Y cells, both apoptotic and necroptotic pathways were activated, as indicated by increased caspase-3 (aCasp-3) activity and phosphorylated MLKL (p-MLKL), respectively (**Fig. 6H-I**). Importantly, AcNPs treatment reduced p-MLKL levels without reducing caspase-3 activity (**Fig. 6H-I**), indicating that lysosomal re-acidification preferentially attenuates TNF-induced necroptosis rather than apoptosis. To further examine the potential involvement of lysosomes in this response, we assessed the spatial relationship between lysosomes and cell-death markers by immunofluorescence. Consistent with the reported association between lysosomal membrane damage and necroptosis, in which activated MLKL can localize to lysosomal membranes and promote LMP [58], we observed substantial colocalization of lysosomes with p-MLKL puncta in TNF-treated cells (**Fig. 6J**), whereas little colocalization was observed with cleaved caspase-3 (**Fig. 6K**). AcNPs treatment markedly reduced p-MLKL-associated lysosomal puncta, concomitant with reduced LMP. These findings raise the possibility that lysosomal degradation of p-MLKL contributes to the attenuation of TNF-induced necroptotic signaling following AcNPs-mediated lysosomal re-acidification.

### Stereotaxic injection of AcNPs restore autolysosomal functions and attenuate necroptosis in the hippocampal region of the APP^NL-G-F^ mouse model of AD

Having observed the effects of TNF in autolysosomal dysfunction and necroptosis in SH-SY5Y cells, we sought to investigate whether similar pathogenic mechanisms are present in the APP^NL-G-F^ mouse model of AD and whether AcNPs can provide therapeutic effect in vivo. Glial activation and neuroinflammation are known to be present in the APP^NL-G-F^ mice starting at 2-month-old [59], which has been shown to peak at 4-month-old with a sustained but lower level of inflammation at a later age [60]. We selected an early age of WT and APP^NL-G-F^ mice at 3-month-old with glial activation (**Fig. S6A**) and elevated TNF level (**Fig. S6B-C**) for stereotaxic injection of PBS (control) and AcNPs into the mouse hippocampus and sacrificed the mice at 4-month-old.

We first showed that there is a significant increase in TNFR1 and p-MLKL levels with no change in the total MLKL level in APP^NL-G-F^ mouse brains as compared to the WT controls (**Fig. 7A-B**), indicating the presence of necroptotic process. We also illustrated that there is no significant change in caspase activities in APP^NL-G-F^ mouse brains (**Fig. 7A-B**). We then observed similar reduction in lysosomal V-ATPase subunits (ATP6V1A, ATP6V1H, and ATP6V1C1) and accumulation of p62, indicating autolysosomal impairment in APP^NL-G-F^ mouse brains as compared to the WT controls (**Fig. 7A-B**). Importantly, we showed that necroptosis activation and autolysosomal impairments are present in the hippocampal CA1 region of APP^NL-G-F^ mice (**Fig. 7C-D and S7A-B**). This makes APP^NL-G-F^ mice a good model for our investigation of the effect of AcNPs in restoring lysosomal acidification and attenuating neuronal necroptosis in vivo.

**Fig. 7.**
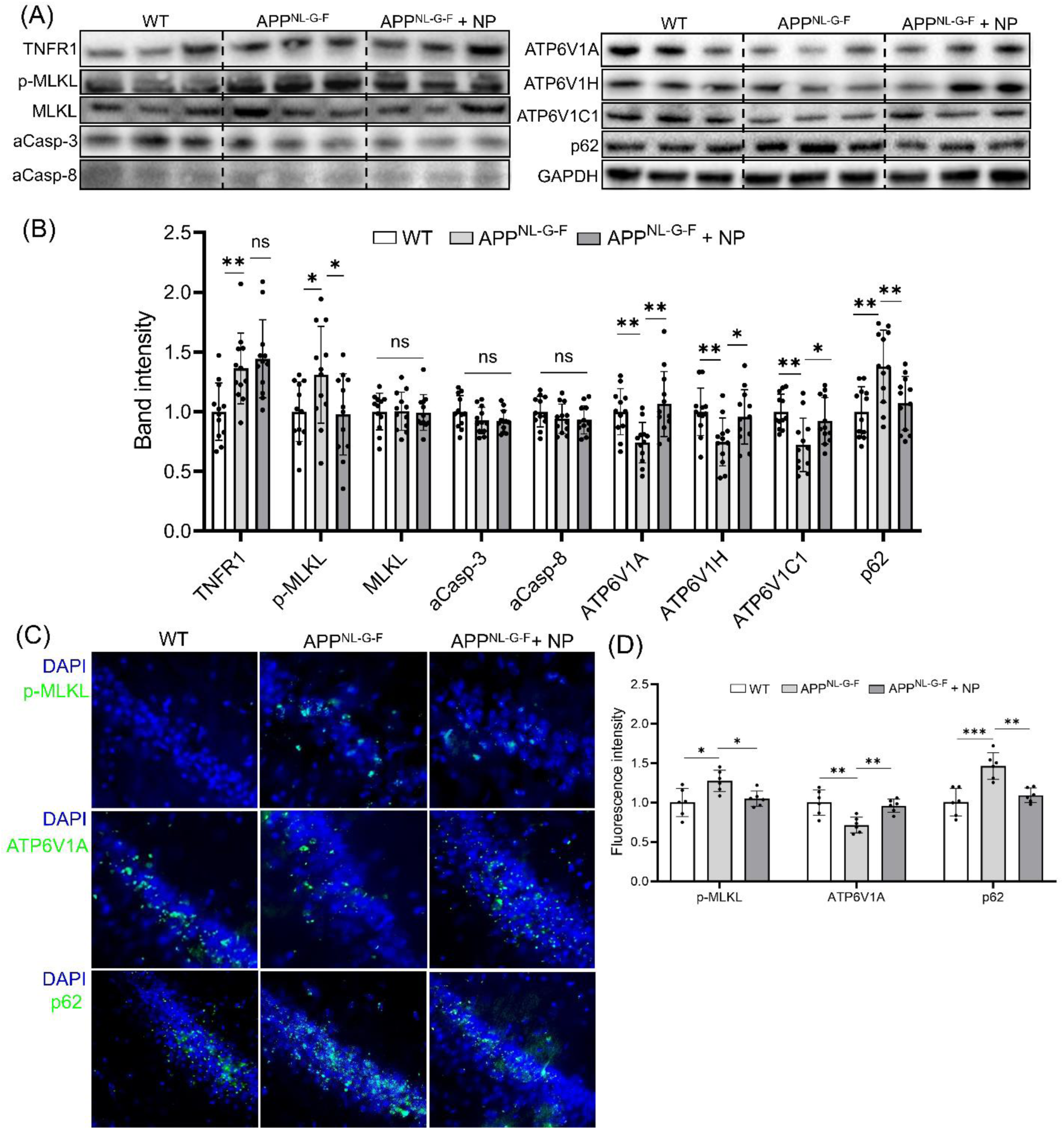
Lysosome-acidifying nanoparticles (AcNPs) restore autolysosomal functions and attenuate necroptotic process in hippocampal region of APP^NL-G-F^ mice. (A-B) Western blotting analysis and quantification for TNFR1 mediated necroptosis (TNFR1, p-MLKL, and MLKL), caspase activities (aCasp-3 and aCasp-8), V-ATPase subunits (ATP6V1A, ATP6V1H, and ATP6V1C1), autophagy receptor p62, and GAPDH as a loading control. (C-D) Immunofluorescence images of p-MLKL, ATP6V1A, and p62 staining in the hippocampal CA1 region of mouse brains of WT, APP^NL-G-F^ and APP^NL-G-F^ injected with AcNPs. The amount of AcNPs (NP) injected is 150 µg per mouse brain hemisphere. Data presented are relative to WT control. Data are means ± SD of N=6-12 mice per group. \**P* < 0.05, \*\**P* < 0.01, \*\*\**P* < 0.001, and ns indicates non-significance by one-way ANOVA with post hoc Tukey’s test for multiple comparisons.

Injection of AcNPs significantly reduced p-MLKL level without augmenting the TNFR1 level, indicating that the attenuation of necroptotic process was not due to reduced TNFR1 expression (**Fig. 7A-B**). We also observed restoration of lysosomal and autophagic functions as shown by an increase in V-ATPase subunits and a decrease in the accumulated p62 upon treatment of AcNPs, respectively (**Fig. 7A-B**). The rescue of autolysosomal dysfunction and attenuation of the necroptotic process were recapitulated in the hippocampal CA1 region (**Fig. 7C-D and S7A-B**). Altogether, our results demonstrated that autolysosomal impairment, in particular lysosomal acidification dysfunction, plays a key role in TNFR1 mediated neuronal necroptosis. Restoration of lysosomal acidification and function may provide an effective approach to attenuate neuronal necroptotic process in AD.

## Discussion

Recent evidence has shown that the activation of neuroinflammatory processes is an early event of AD pathology that occurs due to multiple contributing factors including the accumulation of toxic protein aggregates and glial activation. While toxic protein aggregates may directly induce neuronal death [61,62], recent studies have proposed that the inflammatory responses induced by these toxic protein aggregates may play a leading role in the initiation and subsequent progression of AD [63]. Hyperphosphorylated tau has been shown to induce necroptosis by promoting the necrosome formation and stimulating cell-autonomous cytokine overexpression to propagate inflammation [23]. Aβ accumulation has been shown to lead to necroptosis in AD via microglia activation and cytokine production [25]. Furthermore, Aβ can act as a DAMP to increase the expression of proinflammatory cytokines [64] and elevated cytokine production can further lead to increased toxic protein expression and aggregation which results in an autocatalytic process that propagates the inflammatory processes [65]. Hence, targeting TNFR1-induced necroptosis by promoting autolysosomal degradation represents an important strategy in ameliorating the accumulation of toxic necrosome molecules and protein aggregates [65].

APP^NL-G-F^ mice have been developed to recapitulate expression profiles of risk factor genes, key aspects of neuroinflammation, and Aβ plaque deposition, similar to that in AD patients [59]. With glial activation starting at 2 months of age, it has been suggested that intense neuroinflammation happens between 3-6 months of age in APP^NL-G-F^ with a peak at around 4 months of age [60]. There is corresponding synaptic dysfunction without neuronal death, which is consistent with our observation in the initiation of a low level of necroptotic process. LPS treatment has been shown to increase TNF cytokine level in the hippocampus, accompanied with neuronal death indicated by increased TUNEL staining in APP^NL-G-F^ compared to WT mice at 5-6 months of age [66]. APP^NL-G-F^ mice have also demonstrated early metabolic defects starting from 2 months of age, such as mitochondrial dysfunction, autophagic impairment, and lysosomal defects [67–69]. Downregulation of lysosomal V-ATPase subunits has also been shown in APP^NL-G-F^ [68] and human AD brains [70–72], consistent with our results and the notion of lysosomal impairment as the terminal failure in the autophagic processes.

Although sustained inflammation is present [73], a study has shown that there is no significant increase in the neuronal necroptotic process in APP^NL-G-F^ mice at 12 months of age [74]. This could be due to a lower level of inflammation beyond the early ages. Another possibility is that the presence of necroptotic process observed in some studies could be due to externally induced inflammation such as exposure to inflammatory insults [66] and stereotaxic injection procedures [75] that may trigger further neuroinflammatory signaling and exacerbate neuronal impairment. Although not shown in brain cells, it is worth noting that a study has demonstrated that necroptotic cells show reduced release of conventionally secreted cytokines, although the levels of non-conventionally released cytokines were unaffected or elevated due to membrane permeabilization [76]. On the other hand, the presence of necroptosis is widely demonstrated in other rodent models, including APP^NL-G-F^ rats [74] as well as APP/PS1 [24] and 5xFAD [20] mice. This warrants further investigations on the detailed molecular mechanisms in age-dependent cytokine regulation and neuronal dysfunction in different AD mouse models.

Recently, we have described a tripartite pathogenic mechanism of autophagy-lysosome-mitochondria axis in TNFR1 mediated neuronal necroptosis, with lysosome being the central mediator of this pathogenic axis [77]. While previous studies have focused on promoting autolysosome fusion by increasing autophagosome formation [24], we adopted a novel approach to target and remedy lysosomal acidification dysfunction and LMP to improve the autophagic processes. In the present study, TNF induced multiple dimensions of lysosomal dysfunction, including de-acidification, reduced cathepsin activity, altered lysosomal trafficking and membrane dynamics, and LMP. These findings expand our previous understanding of TNF-induced lysosomal dysfunction by demonstrating that inflammatory stress affects not only lysosomal luminal acidity but also the membrane integrity and dynamic processes required for lysosomal function. Lysosomal membrane integrity is closely linked to cellular survival. It has been suggested that limited release of proteases to the cytoplasm induces apoptosis, while massive LMP results in acute necroptosis [78], indicating that susceptibility of lysosomes to rupture is a key determinant for the propensity of occurrence of necroptotic process [79]. In response to lysosomal membrane injury, the endosomal sorting complex required for transport (ESCRT) machinery has been reported to repair lysosomal membrane and preserve lysosomal acidification [47], while persistent damage can activate lysophagy and clearance of irreparably damaged lysosomes [46,47]. Consistent with this model, we observed increased CHMP2B and TRIM16 following TNF stimulation, suggesting activation of lysosomal membrane damage and quality-control responses. AcNPs treatment reduced both responses together with LMP, suggesting that restoration of lysosomal acidification reduces the underlying lysosomal stress and consequently the requirement for activation of these pathways. Given the established role of LAPTM4B in lysosomal membrane stability and autophagic function, its reduction under TNF stimulation may contribute to the loss of lysosomal membrane homeostasis. We also observed reduced levels of Arl8b and VAMP7 following TNF stimulation, indicating disruption of lysosomal positioning, trafficking, and membrane fusion. Restoration of these proteins following AcNPs treatment further suggests that lysosomal re-acidification helps preserve broader lysosomal membrane homeostasis.

The relationship between lysosomal dysfunction and necroptosis may be particularly important in TNFR1-mediated neuronal injury. We observed that TNF activated both apoptotic and necroptotic pathways, whereas AcNPs treatment preferentially reduced p-MLKL without reducing caspase-3 activity. Moreover, p-MLKL puncta showed substantial colocalization with lysosomes, whereas cleaved caspase-3 showed limited lysosomal colocalization. Recent studies have demonstrated that activated MLKL can localize to lysosomal membranes and contribute to LMP during necroptosis [58], providing a potential mechanistic link between lysosomal membrane damage and necroptotic signaling. Our observation that AcNPs treatment reduced lysosome-associated p-MLKL puncta raises the possibility that restoration of lysosomal degradation facilitates p-MLKL clearance and thereby limits TNF-induced necroptotic signaling. Although direct lysosomal degradation of p-MLKL remains to be established, these findings support functional crosstalk between lysosomal membrane homeostasis and the necroptotic pathway.

Importantly, our findings further connect lysosomal dysfunction to mitochondrial abnormalities. Defective autolysosomal degradation under TNF stimulation was accompanied by impaired mitochondrial turnover, mitochondrial fragmentation, loss of mitochondrial membrane potential, and increased ROS production. Restoration of lysosomal acidification by AcNPs improved autophagic clearance and mitochondrial turnover, accompanied by restoration of mitochondrial morphology and membrane potential and reduction of mitochondrial ROS. These observations are consistent with previous reports showing that LMP related cell death is coordinated in a mitochondria-dependent manner and that p-MLKL can translocate to mitochondria to induce impairments [80], including decreased MMP and increased ROS production [17,32,81]. This supports a feed-forward interaction between lysosomal and mitochondrial dysfunction. Thus, restoring lysosomal acidification may interrupt this pathogenic cycle at an upstream point by improving autophagic degradation and mitochondrial quality control.

Both neuroinflammation [2,3] and lysosomal acidification defect [82] have been described as early pathologies of AD. Our findings suggest that these processes are functionally linked, with lysosomal acidification serving as a key point of intervention in the TNFR1-autophagy-lysosome-mitochondria-necroptosis pathway. Notably, AcNPs did not rescue LPS-induced cell death, consistent with the absence of a lysosomal acidification defect under LPS stimulation and supporting the specificity of this intervention for lysosomal acidification dysfunction. In APP^NL-G-F^ mice, AcNPs treatment similarly restored lysosomal and autophagic function and reduced necroptotic signaling, providing in vivo support for targeting lysosomal dysfunction in the context of AD-associated neuroinflammation. As TNF signaling can be neurodegenerative and neuroprotective, it is important to understand the contribution of individual type of brain cells to neuroinflammation to determine which inflammatory processes are good or bad under disease condition. Hence, it is important to elucidate the role of TNFR1 mediated necroptosis and lysosomal dysfunction in other cell types such as microglia [83,84] and astrocytes [85,86]. It will also be important to determine whether lysosomal damage-response pathways, including ESCRT-mediated repair and lysophagy, are sufficient to restore lysosomal function or become overwhelmed during prolonged inflammatory stress.

In conclusion, our findings identify lysosomal dysfunction as an important component of TNFR1-mediated neuronal necroptosis in AD. TNF-induced lysosomal de-acidification is accompanied by impaired proteolytic function, disrupted membrane dynamics, LMP, impaired autophagic and mitochondrial turnover, and mitochondrial dysfunction, contributing to necroptotic cell death. Restoration of lysosomal acidification with AcNPs reverses multiple defects across this pathway and preferentially attenuates necroptotic signaling. With the notion of lysosome being a cellular signaling hub [87], we propose that lysosome-targeting therapies [88,89] may provide new therapeutic directions for treatment of AD and other neuroinflammatory diseases.

## Authors’ contributions

C.H.L. conceived and directed the research project. J.Z. conducted the cell and animal experiments. K.A. assisted with cell experiments, immunostaining, and image acquisition. G.W.Z.L. assisted with animal studies, immunostaining, and image acquisition. E.N.S. provided support with cell culture and biochemical assays. L.M.O. and J.I. assisted with bioinformatics and image analysis. R.R. and A.M.B. provided critical comments and edited the manuscript. J.Z. and C.H.L. wrote the manuscript. All authors approved the final version of the manuscript.

## Acknowledgements

The authors thank Jia Hui Wong, Enoch Kwok, and Evridiki Asimakidou for their technical support and discussion in this study. The authors also thank the funding sources for supporting this work.

## Supplementary Figures

**Figure S1.**
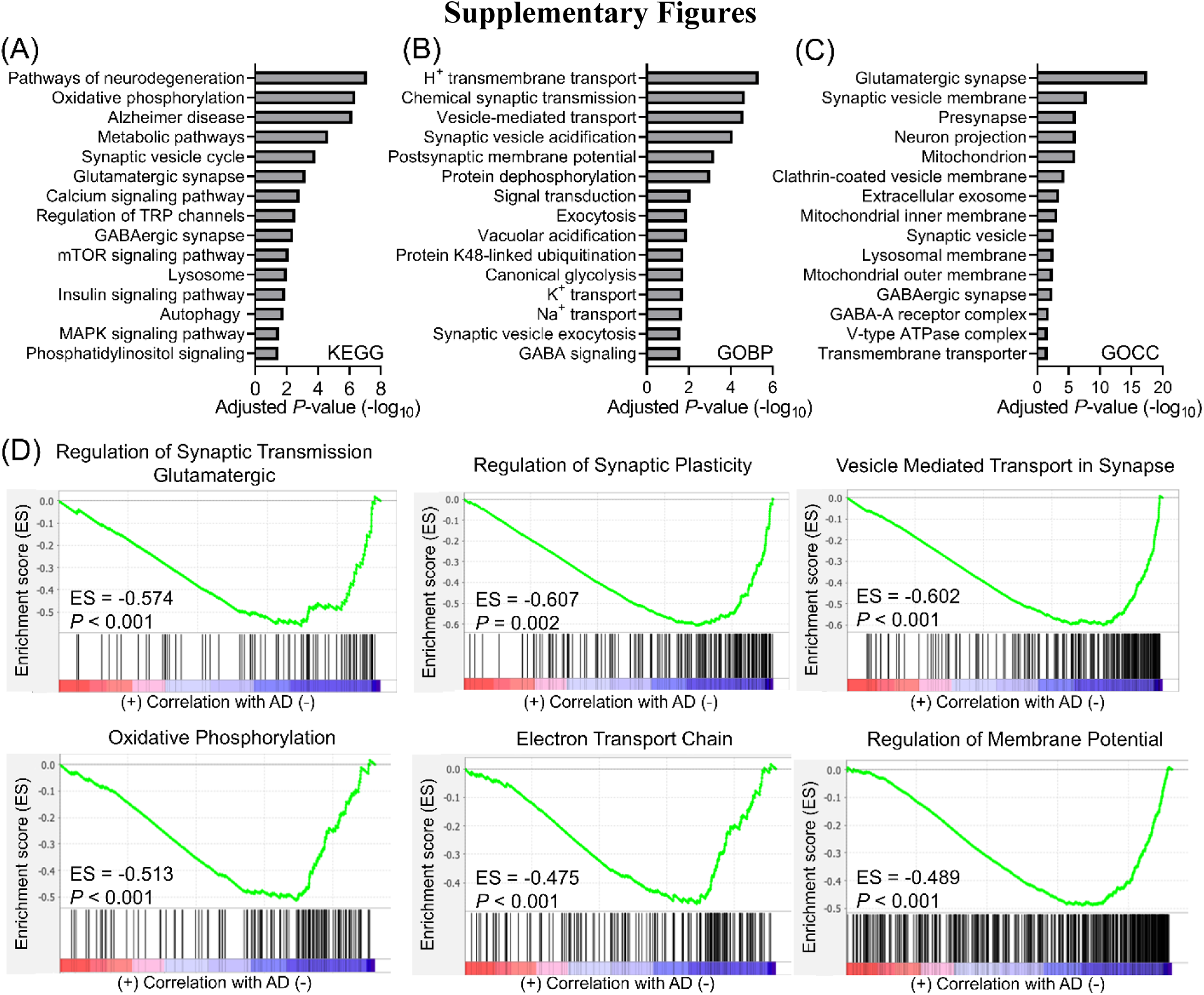
Pathway enrichment analysis and GSEA analysis of downregulated DEGs. (A-C) Pathway enrichment analysis of downregulated DEGs in the GSE173955 dataset using (A) KEGG, (B) GOBP, and (C) GOCC databases. (D) Additional GSEA analysis of the GSE173955 dataset illustrating downregulation of synaptic and mitochondrial functions.

**Figure S2.**
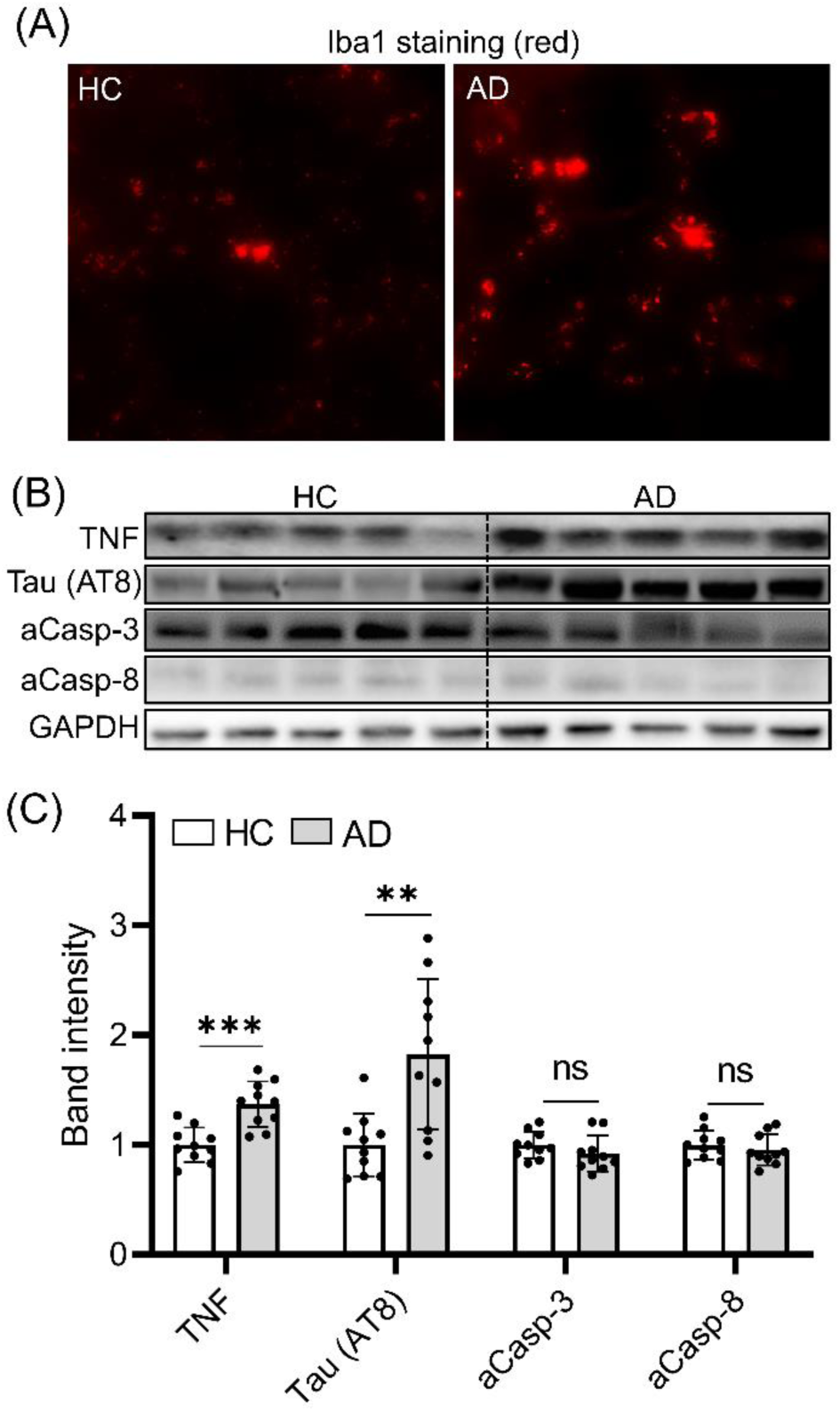
Microglia activation and elevated TNF level in human AD brains. (A) Immunofluorescence images of HC and AD brains stained with Iba1 to determine the extent of microglia activation. (B-C) Western blotting analysis and quantification of the expression level of TNF, Tau (AT8), and caspase activities (aCasp-3 and aCasp-8) in human HC and AD brains with GAPDH as a loading control. Data presented are relative to HC. Data are means ± SD of N=10 HC and N=10 AD brains. \*\**P* < 0.01 and \*\*\**P* < 0.001, and ns indicates non-significance by unpaired Student’s t test for comparison between two samples.

**Figure S3.**
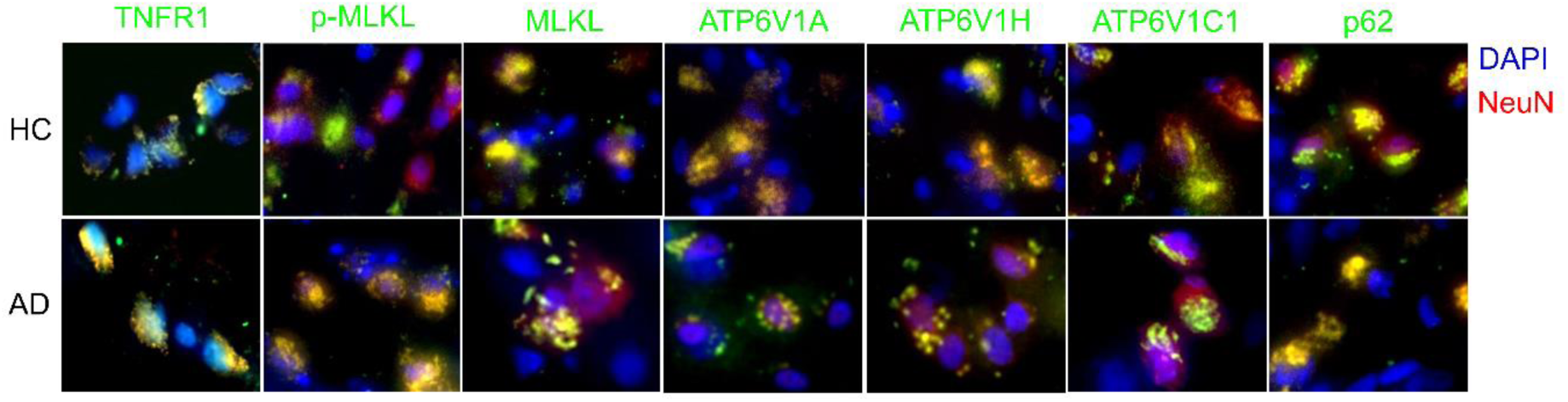
Immunofluorescence staining of human brain tissues with respective markers. Immunofluorescence images showing the presence of TNFR1, p-MLKL, MLKL, V-ATPase subunits (ATP6V1A, ATP6V1H, and ATP6V1C1), and autophagy receptor p62 in hippocampal neurons of HC and AD brain tissues.

**Figure S4.**
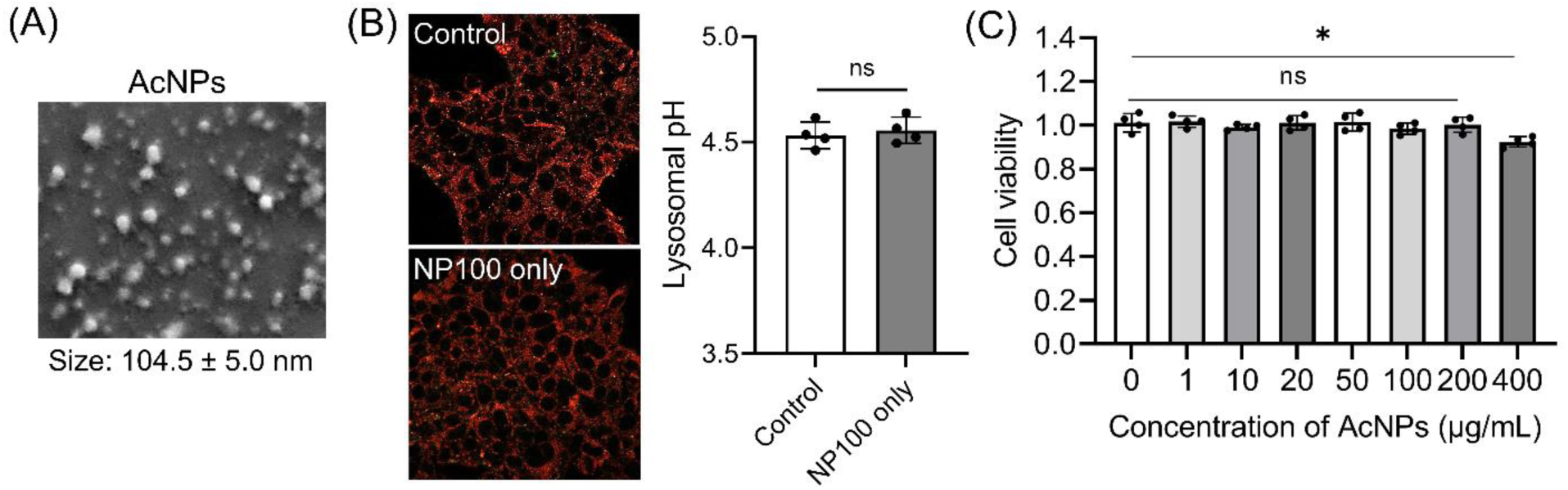
AcNPs are uniform in size and non-toxic to SH-SY5Y cells. (A) Characterizations of AcNPs using SEM and DLS to determine their morphology and size. (B) Lysosomal pH measurement and quantification with LysoSensor Yellow/Blue dye in SH-SY5Y control cells with and without addition of AcNPs. (C) Measurement of cell viability in SH-SY5Y control cells with increasing doses of AcNPs treatment (1 to 400 µg/mL). Data presented are relative to control cells without AcNPs treatment. Data are means ± SD of N=4 independent experiments. NP100 indicates AcNPs treatment at 100 µg/mL. \**P* < 0.05 and ns indicates non-significance by unpaired Student’s t test for comparison between two samples and one-way ANOVA with post hoc Tukey’s test for multiple comparisons.

**Figure S5.**
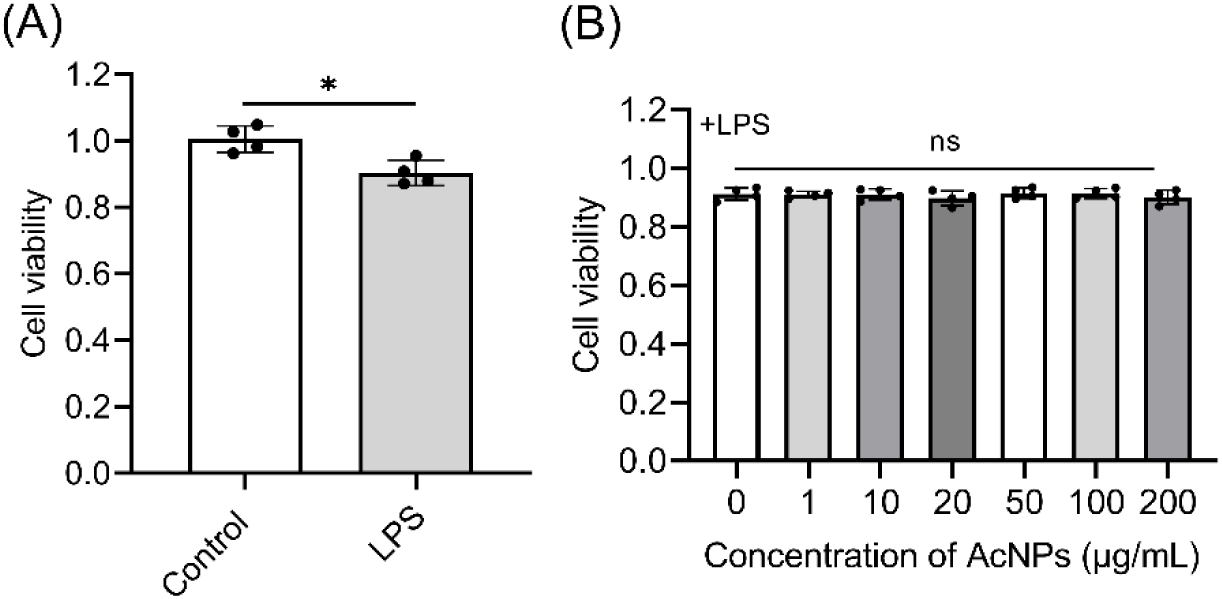
AcNPs do not rescue LPS induced cell death in SH-SY5Y cells. (A) Measurement of cell viability of SH-SY5Y cells with and without LPS (1 µg/mL) treatment. (B) Measurement of cell viability of LPS-treated SH-SY5Y cells with addition of increasing doses of AcNPs (1 to 200 µg/mL). AcNPs do not rescue LPS induced cell death in SH-SY5Y cells. Data presented are relative to control without LPS and AcNPs treatment. Data are means ± SD of N=4 independent experiments. \**P* < 0.05 and ns indicates non-significance by unpaired Student’s t test for comparison between two samples and one-way ANOVA with post hoc Tukey’s test for multiple comparisons.

**Figure S6.**
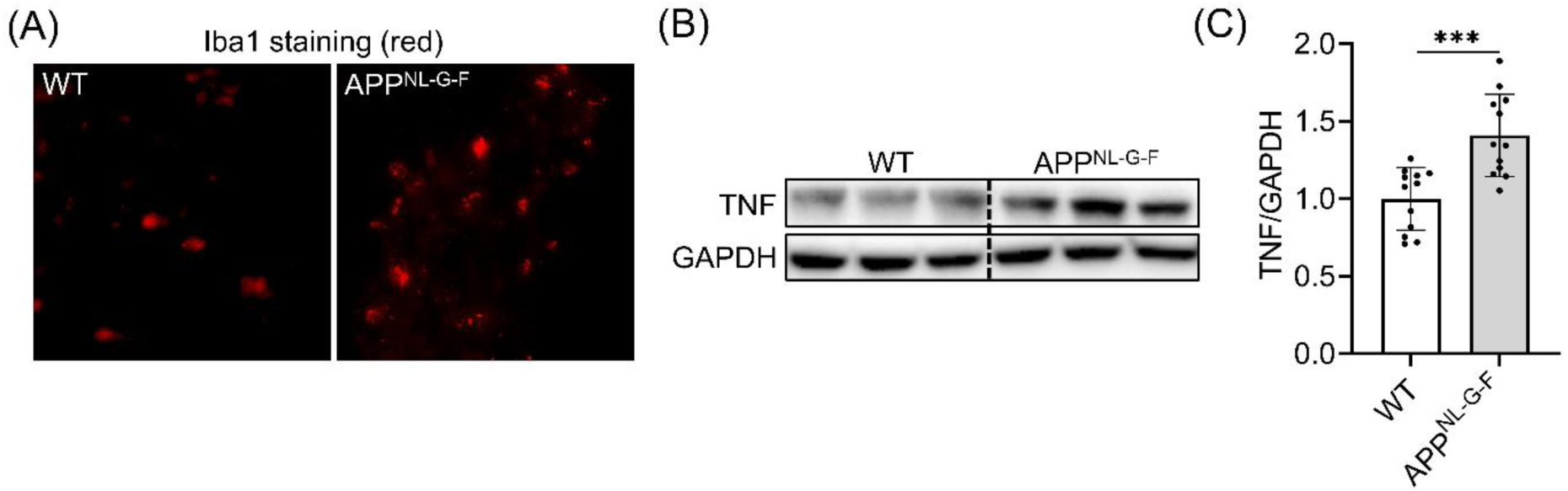
Microglia activation and elevated TNF level in APP^NL-G-F^ mouse brains. (A) Immunofluorescence images of WT and APP^NL-G-F^ mouse brains stained with Iba1 to determine the extent of microglia activation. (B-C) Western blotting analysis and quantification of the expression level of TNF in WT and APP^NL-G-F^ mouse brains with GAPDH as a loading control. Data presented are relative to WT control. Data are means ± SD of N=12 mice per group. \*\*\**P* < 0.001 by unpaired Student’s t test for comparison between two samples.

**Figure S7.**
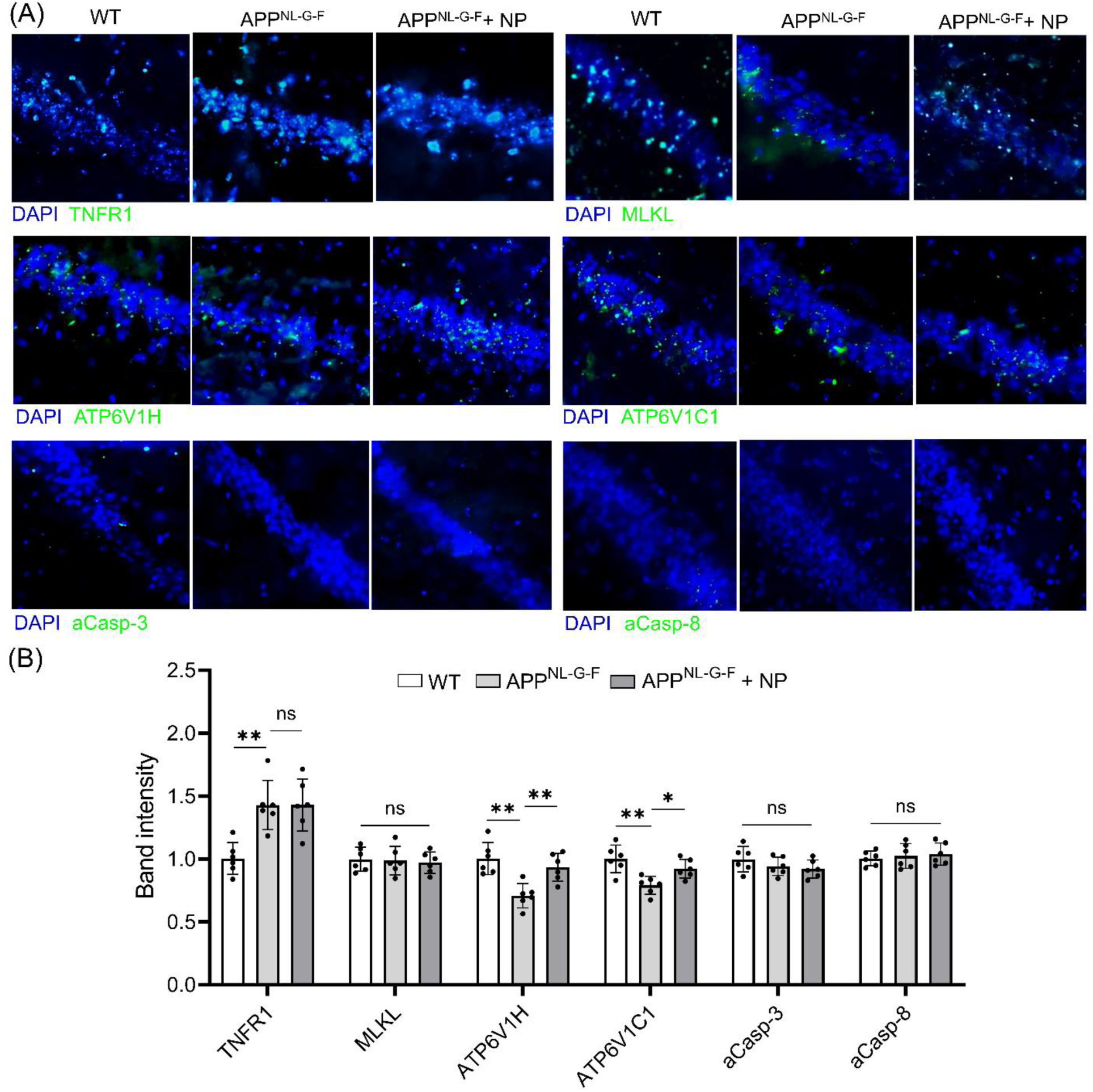
Immunofluorescence staining of mouse brain tissues with respective markers. Immunofluorescence images showing the presence of TNFR1, MLKL, and V-ATPase subunits (ATP6V1H, and ATP6V1C1) as well as caspase activities (aCasp-3 and aCasp-8) in the hippocampal CA1 region of mouse brains under respective conditions. The amount of AcNPs (NP) injected is 150 µg per mouse brain hemisphere. Data presented are relative to WT control. Data are means ± SD of N=6 mice per group. \**P* < 0.05, \*\**P* < 0.01, and ns indicates non-significance by one-way ANOVA with post hoc Tukey’s test for multiple comparisons.

**Table S1.** Demographic, neuropathological, and clinical characteristics of the study cohort.

| **Case** | **Sex** | **Age** | **Braak stage** | **Clinical diagnosis** |
| --- | --- | --- | --- | --- |
| HC1 | M | 85 | II | Non-AD control |
| HC2 | M | 83 | III | Non-AD control |
| HC3 | M | 78 | II | Non-AD control |
| HC4 | M | 93 | II | Non-AD control |
| HC5 | M | 92 | II | Non-AD control |
| HC6 | M | 75 | II | Non-AD control |
| HC7 | F | 74 | I | Non-AD control |
| HC8 | F | 95 | II | Non-AD control |
| HC9 | F | 89 | II | Non-AD control |
| HC10 | F | 78 | III | Non-AD control |
| AD1 | M | 86 | IV | AD |
| AD2 | M | 80 | VI | AD |
| AD3 | M | 80 | V | AD |
| AD4 | M | 86 | V | AD |
| AD5 | M | 87 | V | AD |
| AD6 | F | 66 | VI | AD |
| AD7 | F | 91 | V | AD |
| AD8 | F | 93 | V | AD |
| AD9 | F | 87 | V | AD |
| AD10 | F | 73 | VI | AD |

**Table S2.** List of primary and secondary antibodies used in this study.

| **Antibodies** | **Source** | **Identifier** | **Application** |
| --- | --- | --- | --- |
| Anti-TNFR1 | Proteintech | 60192-1-Ig | IF and WB |
| Anti-ATP6V1A | Proteintech | 17115-1-AP | IF and WB |
| Anti-ATP6V1H | Proteintech | 26683-1-AP | IF and WB |
| Anti-ATP6V1C1 | Proteintech | 16054-1-AP | IF and WB |
| Anti-MLKL | Proteintech | 21066-1-AP | IF and WB |
| Anti-Galectin-3 | Proteintech | 14979-1-AP | IF |
| Anti-p-MLKL (Ms) | Abcam | ab196436 | IF and WB |
| Anti-p-MLKL (Hu) | Abcam | ab187091 | IF and WB |
| Anti-TNF | Abcam | ab6671 | WB |
| Anti-SQSTM/p62 | Cell Signaling Technology | 5114 | IF and WB |
| Anti-LC3A/B | Cell Signaling Technology | 12741 | WB |
| Anti-GAPDH | Cell Signaling Technology | 2118 | WB |
| Anti-Rabbit (HRP) | Cell Signaling Technology | 7074 | WB |
| Anti-Mouse (HRP) | Cell Signaling Technology | 7076 | WB |
| Anti-Cleaved Caspase-3 | Cell Signaling Technology | 9661 | IF and WB |
| Anti-Cleaved Casapase-8 | Novus Biologicals | NB100-56116 | IF and WB |
| Anti-LAPTM4B | Proteintech | 18895-1-AP | IF |
| Anti-TRIM16 | Thermofisher Scientific | PA5-104489 | IF |
| Anti-VAMP7 | Biotechne | NBP1-07118 | IF |
| Anti-Arl8b | Proteintech | 13049-1-AP | IF |
| Anti-CHMP2B | Thermofisher Scientific | PA5-31128 | IF |
| Anti-NeuN | Synaptic Systems | 266004 | IF |
| Anti-Iba1 | Synaptic Systems | 234009 | IF |
| Anti-Phospho-Tau (AT8) | Thermofisher Scientific | MN1020 | WB |
| Goat anti-Rabbit (AF488) | Thermofisher Scientific | A-11008 | IF |
| Goat anti-Mouse (AF488) | Thermofisher Scientific | A-11001 | IF |
| Goat anti-Rabbit (AF594) | Thermofisher Scientific | A-11012 | IF |
| Goat anti-Mouse (AF594) | Thermofisher Scientific | A-32742 | IF |
| Goat anti-Guinea Pig (AF594) | Thermofisher Scientific | A-11076 | IF |
| Goat anti-Chicken (AF594) | Thermofisher Scientific | A-11042 | IF |
| DAPI | Thermofisher Scientific | A-9542 | IF |

